# Transcriptomic correlates of state modulation in GABAergic interneurons: A cross-species analysis

**DOI:** 10.1101/2023.12.04.569849

**Authors:** Joram Keijser, Loreen Hertäg, Henning Sprekeler

## Abstract

GABAergic inhibitory interneurons comprise many subtypes that differ in their molecular, anatomical and functional properties. In mouse visual cortex, they also differ in their modulation with an animal’s behavioural state, and this state modulation can be predicted from the first principal component (PC) of the gene expression matrix. Here, we ask whether this link between transcriptome and state-dependent processing generalises across species. To this end, we analysed seven single-cell and single-nucleus RNA sequencing datasets from mouse, human, songbird, and turtle forebrains. Despite homology at the level of cell types, we found clear differences between transcriptomic PCs, with greater dissimilarities between evolutionarily distant species. These dissimilarities arise from two factors: divergence in gene expression within homologous cell types and divergence in cell type abundance. We also compare the expression of cholinergic receptors, which are thought to causally link transcriptome and state modulation. Several cholinergic receptors predictive of state modulation in mouse interneurons are differentially expressed between species. Circuit modelling and mathematical analyses suggest conditions under which these expression differences could translate into functional differences.

## Introduction

GABAergic inhibitory interneurons are a highly diverse population consisting of multiple cell types [1, 2]. In recent years, single-cell RNA sequencing (scRNA-seq [3]) has revealed that these types can be further subdivided into tens of subtypes [4, 5, 6] that also differ in their morphological and electrophysiological properties [7, 8]. So far, it has been difficult to understand the functional relevance of this fine-grained diversity. Bugeon et al. [9] recently bridged this gap by revealing that interneurons show subtype-specific modulation with an animal’s behavioural state, at least in layers 1-3 of mouse primary visual cortex (VISp). Strikingly, this state modulation could be predicted from the first transcriptomic principal component (tPC1). An interneuron’s tPC1 score also correlated with other dimensions of interneuron diversity, such as electrophysiology and connectivity, hinting at an “approximate but general principle” of mouse cortical interneurons [9].

Intrigued by these findings, we wondered how general the principle embodied by tPC1 actually is (Fig. 1). Are transcriptomic correlates of state modulation similar across different species, or at least across mouse cortical layers and areas? If yes, this similarity would suggest conserved principles; if no, the difference could reveal distinct solutions to shared computational problems [10, 11, 12]. The uniformity of interneurons in the mouse brain [6] suggests that their gene expression and state modulation patterns observed in VISp might apply generally. In fact, earlier work by the authors of ref. [9] found that hippocampal interneurons are also organised along a single latent factor [13]. Similarly, recent comparative transcriptomic analyses have emphasized the conservation of (cortical) inhibitory interneurons across mammals [14, 15, 16, 17], and more distantly related species [18, 19, 20]. But these and other studies [21, 22] have also discovered species-specific interneuron subtypes. Additionally, the relative proportions of interneuron types vary even across mouse cortex [23, 24], as does the modulation of interneurons with brain state [25, 26, 27, 28].

**Figure 1:**
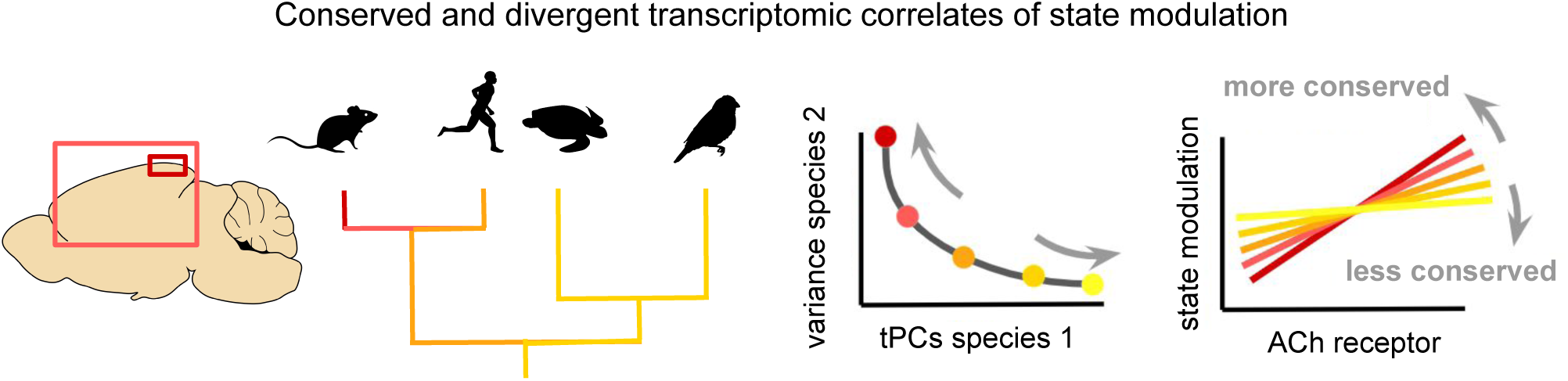
Schematic of our main question: could the same transcriptomic axis predict state modulation in other layers and areas of the mouse cortex and in other species? We investigate this by comparing transcriptomic principal components (tPCs) and cholinergic (ACh) receptor expression across RNA-seq datasets [9, 15, 19, 14, 6, 18, 29].

We therefore investigated the transcriptomic correlates of state modulation in seven existing single-cell RNA-seq (scRNA-seq) and single-nucleus RNA-seq (snRNA-seq) datasets from the forebrains of mice [9, 6, 29], humans [14, 15], turtles [18], and songbirds [19]. These species each have homologous types of inhibitory interneurons, but their evolutionary history and brain organization vary in important ways. Reptilian cortex, for example, has a three-layered structure akin to the mammalian piriform cortex [30, 31]. Yet, transcription factor expression rules out a one-to-one homology between individual reptilian and mammalian layers or projection cell types [32, 18, 33]. The songbird forebrain, on the other hand, is organised in nuclei instead of layers [34, 35, 36]. We found that transcriptomic PCs show relatively minor differences across smaller evolutionary distances (e.g., between mice and humans) but diverge over longer evolutionary time scales (e.g., mice and turtles). Between-species differences dwarf within-species differences, likely due to biological rather than technical reasons. Specifically, we trace differences in tPCs to species-specific cell type abundances and within-type gene expression patterns. We also find a combination of conservation and divergence in the expression of the cholinergic receptors correlated with state modulation in mice [9]. Circuit modelling predicts the connectivity patterns for which differences in receptor expression translate into species-specific state modulation of interneurons and cortical information flow.

## Results

We first validated our analysis pipeline by replicating the relevant results from Bugeon et al. [9] on their data and conducting several additional analyses. Briefly, we reproduced the systematic variation of interneuron subtypes with behavioural state (roughly, running vs stationary) and its correlation with tPC1 (Fig. S1). This correlation seems driven by differences within and across cell types [37] and is strongest within the Pvalb and Sst populations (Fig. S2). Whether interneurons form a continuum or cluster along tPC1 depends on the preprocessing of the transcriptomic data (Fig. S3). These caveats aside, our analyses are consistent with those from Bugeon et al. and might reveal similar patterns—or the lack thereof— in other brain areas and species. A detailed description of the replication can be found in the supplementary material (see Replication of Bugeon et al).

### Conserved and divergent transcriptomic axes across species

Having validated our approach on mouse data, we next turn to the cross-species comparison using transcriptomic data from humans (*Homo sapiens*, [15]), turtles (*Trachemys scripta elegans*, [18]), and zebra finch (*Taeniopygia guttata*, [19]); see Table 1 for an overview of all analysed datasets. We compare these data with a large reference dataset from mouse VISp [6].

**Table 1:** Overview of analysed datasets. Sn: single nucleus, sc: single cell. SS: Smart-Seq. M1: primary motor cortex, VISp: primary visual cortex, HVC: high vocal centre, MTG: middle temporal gyrus, Ctx: cortex, Hpc: hippocampal formation. Genes/cell: median number of genes detected per cell. Reads: median number of reads per cell. For the single nucleus data, reads aligned to exons and introns were used. The data from ref. [9] comprised real-valued estimates of counts (see Methods from [9]; the number of reads was therefore unavailable.

| Name | Species | Area(s) | Technology | Cells | Genes/cell | Reads |
| --- | --- | --- | --- | --- | --- | --- |
| Bugeon [9] | Mus musculus | VISp L1-3 | CoppaFish (sc) | 1,065 | 49 | - |
| Tasic [6] | Mus musculus | VISp | SS v4 (sc) | 6,125 | 9,795 | 2,009,806 |
| Yao [29] | Mus musculus | Ctx & Hpc | 10x V2 (sc) | 177,614 | 3,750 | 8,957 |
| Bakken [15] | Homo sapiens | M1 | SS v4 & 10X (sn) | 23,992 | 4,719 | 14,337.5 |
| Hodge [14] | Homo sapiens | MTG | SS V4 (sn) | 4,164 | 8,344.5 | 1,901,796.5 |
| Colquitt [19] | Taeniopygia guttata | HVC & RA | 10X v2 (sn) | 3,786 | 161.5 | 3,150.5 |
| Tosches [18] | Trachemys scripta elegans | Ctx | Drop-Seq v3 (sc) | 640 | 2,952 | 6,284 |

**Table 2:** Datasets and preprocessing steps for main data analysis figures. CP10K: counts per ten thousand scaling; log: log-transform, one2one: one-to-one orthologs; HVGs: highly variable genes; avg. subtypes: average expression and state modulation within each subtype. downs.: downsample gene counts.

| Figure | Datasets | Preprocessing |
| --- | --- | --- |
| Fig. 2 | [15], [9], [19], [6], [18] | CP10K, log, one2one, HVGs |
| Fig. 3 | [9], [6], [29] | CP10K, log, one2one, HVGs |
| Fig. 4 | [15], [9], [19], [6], [18] | one2one, avg. subtypes, z-score (dot plots) |
| Fig. S1 | [9], [6] | CP10K, log, avg. subtypes |
| Fig. S2 | [9] | CP10K, log |
| Fig. S3 | [9] | CP10K, log (top row) |
| Fig. S4 | [15], [19], [6], [18] | Downs., CP10K, log, one2one, HVGs |
| Fig. S6,S7, S8 | [15], [19], [6], [18] | CP10K, log, one2one, HVGs |
| Fig. S9 | [15], [14], [6], [29] | CP10K, log, one2one, HVGs |
| Fig. S10 | [15],[14] | CP10K, log, one2one, HVGs |
| Fig. S11 | [9], [6] | CP10K, log, avg. subtypes |
| Fig. S12 | [15],[6], [14], [29] | Z-score (dot plots) |

We first visualized the data from different species. To this end, we preprocessed the datasets using the same analysis pipeline and applied PCA to the resulting RNA count matrices (see Materials and Methods). The projection onto the first 2 tPCs of the human, but not turtle or zebra finch data, was similar to that of the mouse data (Fig. 2a). Mouse and human interneurons clustered by developmental area [38], with medial ganglionic eminence (MGE)-born Pvalb and Sst cells occupying one side of tPC1, and caudal ganglionic eminence (CGE)-born Lamp5, Vip, and Sncg cells the other. An intermediate position was occupied by a small group of Meis2 neurons [6], located in the white matter [39]. In contrast to the mammalian datasets, the turtle and finch data were characterised by a large population of Meis2-positive neurons (Fig. 2a, Table 3). Transcriptomic and morphological evidence suggests that these cells are likely homologous to neurons in the mammalian striatum rather than the white matter [18, 19].

**Figure 2:**
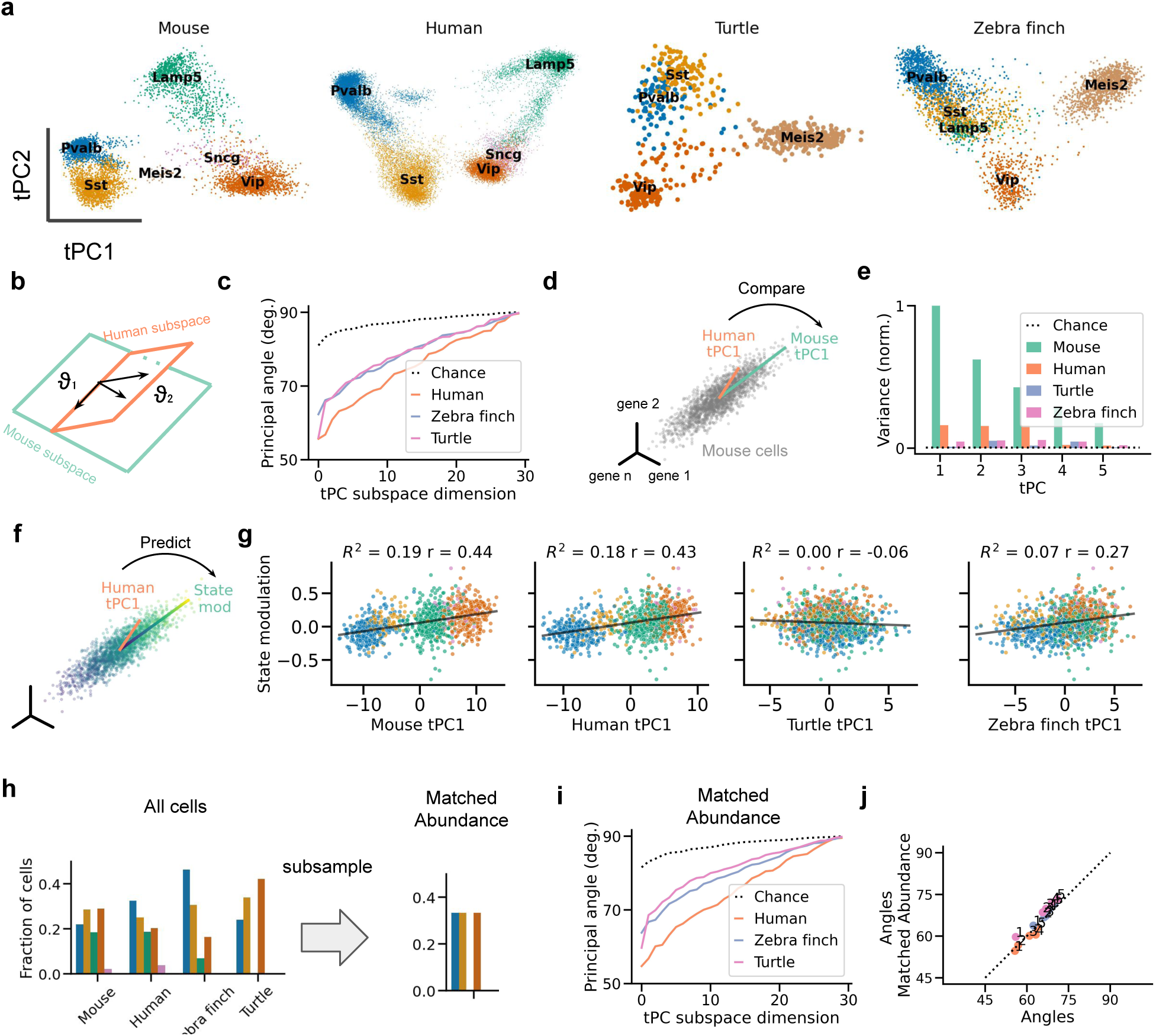
Transcriptomic PCs capture conserved and divergent global gene expression patterns. **(a)** Projections of gene expression data from forebrain interneurons onto each dataset’s first tPCs. Samples correspond to cells; colours indicate cell types. Variance explained by tPC1&2: 23.5% (mouse), 22.4% (human), 13.2% (turtle), and 12.6% (zebra finch). **(b)** Schematic: principal angles measure dissimilarity between subspaces spanned by two sets of tPCs (see Materials and Methods). Larger angles indicate larger differences. **(c)** Principal angles between human and mouse subspaces are smaller than between mouse and zebra finch or turtle subspaces. Chance level estimated by sampling random, normalized vectors. **(d)** Schematic: Variance explained in the mouse data as a measure of tPC similarity. A tPC’s length is proportional to the variance it explains. **(e)** Variance of mouse data explained by tPCs of different datasets, normalized to the variance explained by mouse tPC1. The human, zebra finch, and turtle tPC1 explain 16.2%, 4.6%, and 1.1% of the variance explained by mouse tPC1. A random direction (dashed line) explains 0.5%. **(f)** Schematic: Predicting mouse state modulation from human tPC1. The colour gradient symbolizes the state modulation of mouse cells. **(g)** State modulation of mouse interneurons can be predicted from each interneuron’s projection onto tPC1 from a reference mouse dataset and human tPC1. It can also be predicted, to some extent, from the projection onto zebra finch tPC1, but not turtle tPC1. *R*^2^: cross-validated fraction of variance explained, *r*: Pearson correlation. **(h)** Subsampling procedure to control for the relative abundance of interneuron subclasses across datasets. Colours code for cell types (see (a)). **(i,j)** Matching the relative abundance does not increase the similarity of datasets as measured using principal angles. Numbers indicate order or PCs. Data from refs. [6] (mouse), [9] (mouse state modulation), [15] (human), [18] (turtle), [19] (zebra finch).

**Table 3:** Percentage of cell types for each dataset, rounded to a single decimal place. The mammalian Meis2-positive cells are likely not homologous to the turtle/finch Meis2-positive cells ([19], see text) but are grouped for convenience.

|  | Bakken [15] | Bugeon [9] | Colquitt [19] | Hodge [14] | Tasic [6] | Tosches [18] | Yao [29] |
| --- | --- | --- | --- | --- | --- | --- | --- |
| Pvalb | 32.4 | 27.9 | 31.9 | 17.5 | 21.8 | 13.8 | 17.2 |
| Sst | 25.0 | 8.5 | 21.0 | 30.6 | 28.4 | 19.5 | 26.7 |
| Lamp5 | 18.6 | 35.8 | 4.7 | 27.8 | 18.3 | 0.0 | 23.7 |
| Vip | 20.2 | 24.4 | 11.2 | 24.1 | 28.7 | 24.2 | 24.6 |
| Sncg | 3.7 | 3.5 | 0.0 | 0.0 | 2.0 | 0.0 | 7.8 |
| Meis2 | 0.0 | 0.0 | 31.3 | 0.0 | 0.7 | 42.5 | 0.0 |

We quantified these visual differences using the principal angles, which generalise the notion of angle between two lines in a plane (Fig. 2b; see Materials and Methods). Here, we computed the angles between subspaces spanned by each dataset’s top 30 PCs. Consistent with the impression from the first 2 PCs, the principal angles were the smallest between mouse and human subspaces (Fig. 2c). Turtle and zebra finch PCs were both dissimilar to mouse PCs. Principal angles do not require a one-to-one relationship between individual principal components but also do not consider the variance explained by these components. For instance, a pair of highly similar but low-variance dimensions will result in small principal angles—inadvertently suggesting high similarity. We therefore performed a complementary analysis by computing the variance in the mouse data explained by the PCs of other datasets (Fig. 2d). The first human PC accounted for 16% of the variance explained by the first mouse PC; the turtle and songbird tPC1 accounted for 1% and 5%, respectively (Fig. 2e). Each tPC1 explained more variance than a random direction (0.5%), consistent with some shared global structure.

We confirmed that these results were not due to technical differences in the different datasets. We first controlled for sequencing depth by downsampling gene counts to that of the lowest-depth dataset (Fig. S4, Materials and Methods). We also controlled for sample size by downsampling cells to the number of the smallest dataset (Fig. S5). In either case, the relatively small differences between mouse and human tPCs were maintained, suggesting this effect is not due to technical factors. We also mapped each dataset onto the mouse data using anchor-based integration [40]. This method has been widely used in cross-species analyses (e.g., [14, 19, 41, 42]). As expected, computational integration increased the similarity among the datasets (Fig. S6), but, importantly, it preserved the larger similarity between human and mouse data.

How might the transcriptomic differences relate to state modulation? It is possible that even though the top PCs are only similar between mouse and human interneurons, mouse tPC1 does predict state modulation in other species. Because state modulation information was only available for the mouse [9], we projected this data onto the tPCs from other datasets to determine their predictive ability (Fig. 2f). We validated this approach by showing that tPC1 from a reference mouse dataset predicted state modulation in the Bugeon et al. data (*R*^2^ = 0.19). Next, we found that the human tPC1 predicts state modulation in the mouse approximately as well as mouse tPC1 (*R*^2^ = 0.18), but the turtle tPC1 did not (*R*^2^ = 0) (Fig. 2g; compare with Fig. S1). The zebra finch tPC1 showed a weak but significant ability to predict state modulation (*R*^2^ = 0.07). We conclude that human tPCs are similar to those of the mouse also on a functional level, in line with evolutionary history. We emphasise that this result relies on statistical similarities at the transcriptomic level and will need to be tested using simultaneously recorded neural and behavioural data.

What evolutionary changes underlie the differences between transcriptomic PCs? At least two non-mutually exclusive processes are possible. First, homologous subclasses could evolve in a species-dependent manner, as indicated by differences in gene expression. Second, evolution can also change the relative abundance of otherwise conserved classes [15, 43]. We wondered if the relative abundance of cell classes was sufficient to explain the species differences. To this end, we resampled cells to equal fractions, such that the 3 classes (Pvalb, Sst, Vip) present in all datasets each accounted for one-third of the cells (Fig. 2h). This increased the visual similarity between the first two tPCs of the mammalian and non-mammalian datasets due to the absence of Meis2 neurons (Fig. S7). Still, the matched-abundance datasets were as dissimilar as the original datasets (Figs. 2j, S8). This highlights the divergence of homologous cell types as a driver of evolutionary change in the global transcriptomic landscape.

### Similar transcriptomic axes across mouse datasets

The previous cross-species comparison is based on data collected with different sequencing protocols and from different brain areas. To account for these factors, we calibrated the between-species differences against within-species differences by comparing three mouse datasets (Fig. 3a): the in situ data from VISp layers (L) 1-3 [9], the plate-based (SMART-seq2) data from VISp L1-6 [6], and the droplet-based (10X) data from multiple cortical and hippocampal areas (Ctx & Hpc, [29])

**Figure 3:**
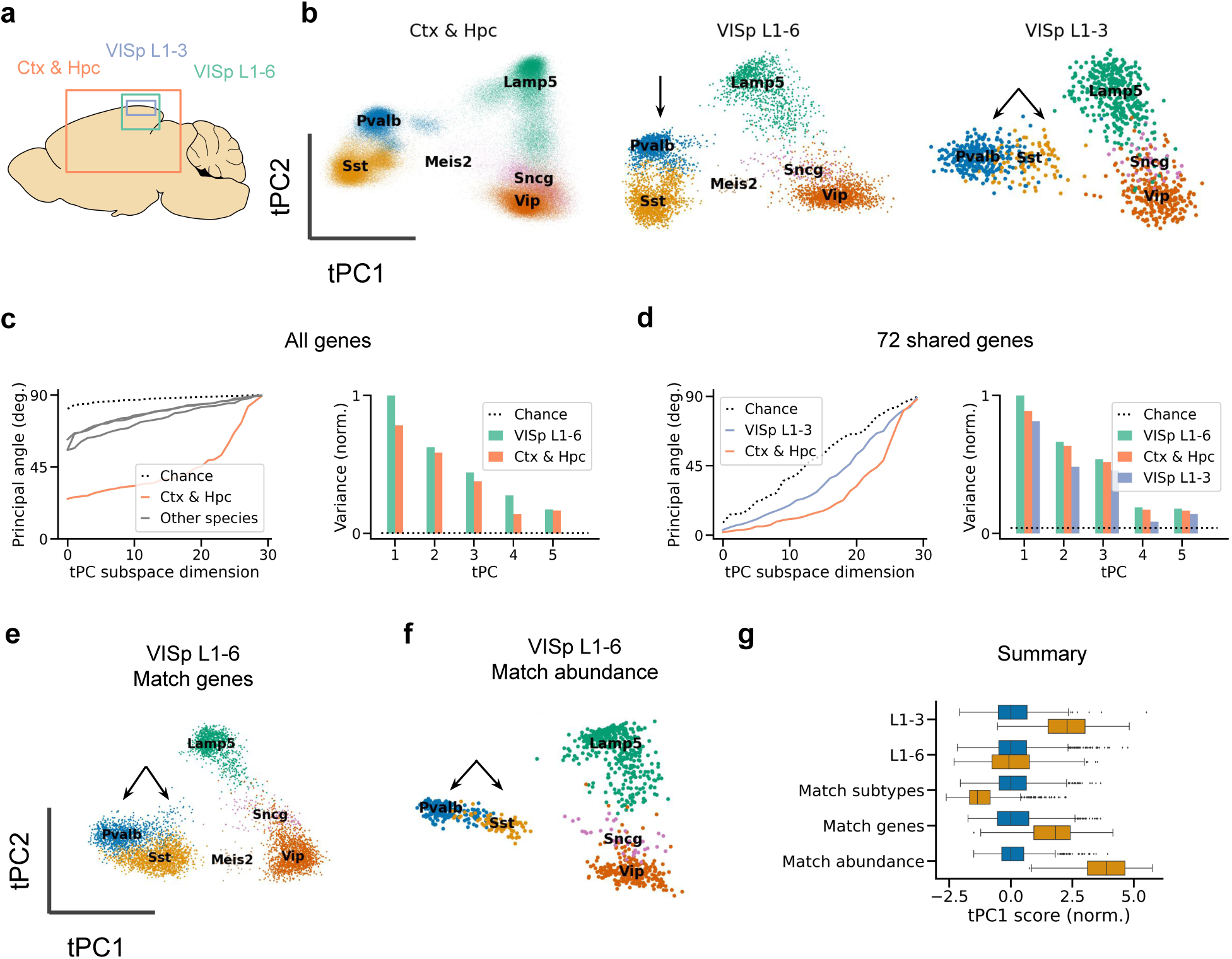
Similar transcriptomic PCs across mouse data sets. **(a)** Schematic of brain areas sequenced for different mouse datasets. Ctx: cortex, Hpc: hippocampal formation, VISp: primary visual cortex. The datasets were also collected using different technologies (Table 1). **(b)** Similar tPC1&2 across mouse datasets; tPCs1&2 jointly explain 17.7%, 23.1%, and 29.8% of variance, from left to right. Arrows indicate a qualitative difference: the relative position of Pvalb and Sst cells along tPC1. **(c)** Quantitative comparison with VISp L1-6 dataset based on 2000 highly variable genes; tPC1 of the Ctx & Hc dataset explains 78.3%% of the variance explained by tPC1 of VISp L1-6. Grey lines: cross-species angles, taken from Fig. 2c. **(d)** As (c), but based on the 72 genes shared by the three datasets. In this reduced space, tPC1 of the Ctx & Hc and VISp L1-3 explains 88.7% and 81.3%, respectively, of the variance explained by VISp L1-6. **(e)** Relatively small change in Pvalb and Sst position after matching gene sets between L1-6 and L1-3 data. **(f)** Larger differences due to relative cell type abundance. **(g)** Distribution of tPC1 projection of Pvalb (blue) and Sst (orange) cells for the L1-3 data and different versions of the L1-6 data. Match subtypes: select only the Sst subtypes present in the L1-3 dataset. Match genes: select only the genes present in the L1-3 dataset. Match abundance: subsample such that Sst cells comprise only 8% of the samples, as in the L1-3 data. Projections were normalized such that the mean and variance of the respective Pvalb population were zero and one, respectively. Expression data from refs. [29] (Ctx & Hpc), [6] (VISp L1-6), and [9] (VISp L1-3).

Visually, the projections onto the first tPCs were similar (Fig. 3b), with interneurons clustering by developmental area, as before. But subtle differences were also visible. For example, the L1-3 dataset lacked a Meis2 population present in both L1-6 datasets (Fig. S9; [6, 29]). The tPC1 score of Sst cells also varied between datasets. In the L1-3 data, Sst cells occupied an intermediate position on tPC1 (and tPC2) compared to Pvalb cells, consistent with their weaker state modulation (Fig. S1b). In contrast, Sst and Pvalb cells occupied similar positions in the other datasets.

The three datasets were also quantitatively similar. Principal angles between different mouse datasets were substantially smaller than angles between species (Fig. 3c). To compare the larger mouse datasets with the smaller dataset of Bugeon et al. [9], we performed the same analyses after selecting the 72 genes shared by all datasets. This revealed the Ctx & Hpc data to be more similar to the VISp L1-6 data than the VISp L1-3 data (Fig. 3d), consistent with the varying relative positions of the cell types in the space of the first two PCs.

Several factors could explain the different positions of Pvalb and Sst cells along tPC1 and tPC2 (arrows in Fig. 3b). We first tested if the difference was due to layer-specific subtypes known to be transcriptomically identifiable (see, e.g., [6, 44]). However, selecting L1-3 subtypes from the L1-6 data followed by PCA only moved the Sst cells further along tPC1 (Fig. 3g, ”match subtypes”). We next tested for the influence of gene set by performing PCA on the L1-6 data after selecting 72 genes describing the L1-3 dataset. This only modestly increased the similarity to the in situ data (Fig. 3e), reflecting the careful selection of the gene panel [45]. Finally, we reasoned that the intermediate position of Sst cells in the L1-3 data could be due to their relative sparsity (8% in the L1-3 data vs 28% in the L1-6 data). After all, a given pattern of covariability explains less variance when present in a smaller number of samples. Indeed, sampling the same number of cells from the entire Sst population moved the Sst population to an intermediate tPC1 position (Fig. 3f). Therefore, the intermediate position of Sst cells in the L1-3 dataset might be due to their relative sparsity.

In summary, mouse datasets are highly similar compared to cross-species datasets despite differences in brain area [6] and sequencing technology [46]. Two human datasets [14, 15] showed equally high levels of similarity (Fig. S10). Between-species differences, therefore, likely reflect biologically meaningful signals rather than technical artefacts.

### Evolution of cholinergic receptor expression

So far, we have shown that interspecies expression differences are reflected in the first principal components. This rules out a conserved tPC1 that predicts state modulation—at least across evolutionarily distant species. However, it does not rule out that the species-specific tPCs predict state modulation. Unfortunately, this cannot be tested directly due to the lack of data on state modulation for the other species. As a proxy, we, therefore, analyzed the expression of cholinergic receptors that are known to contribute to the correlation between tPC1 and state modulation in mice (Fig. 4a, [9]).

**Figure 4:**
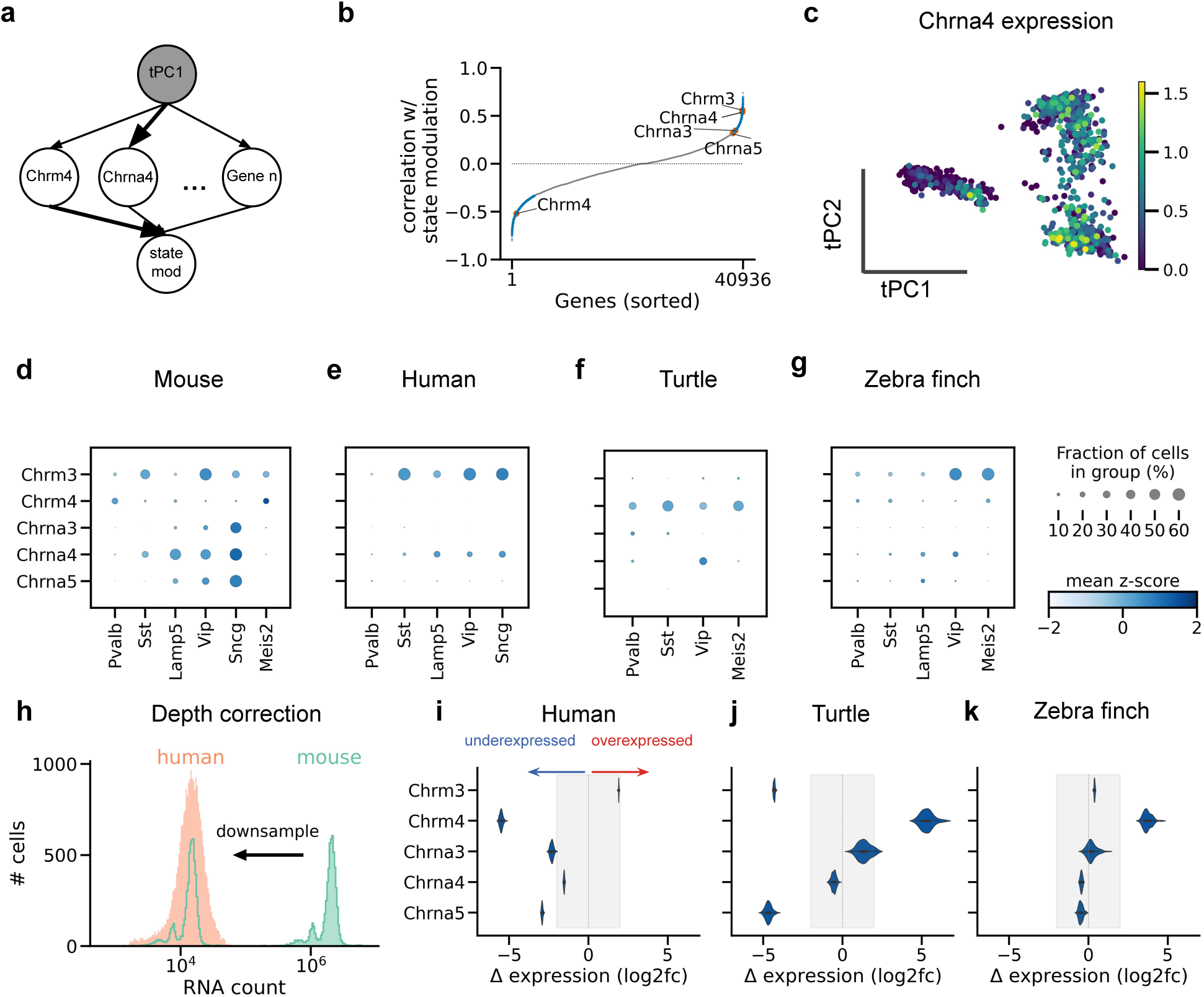
Evolution of cholinergic receptor expression. **(a)** Graphical model of the relationship between tPC1, gene expression, and state modulation. The shaded variable tPC1 is not directly observable. Arrows indicate direct dependence relationships; arrow thickness indicates the strength of the relationship. **(b)** Correlation between gene expression and state modulation in the mouse data. Gene expression and state modulation were not measured in the same cells and were therefore combined at the level of subtypes (Materials and Methods). Blue: significant correlation (*p <* 0.05), grey: not significant. Annotated are the cholinergic receptors that predict state modulation (Fig. S11). Chrm4 is ranked 327th (top 1.5%) with the strongest negative correlation. Chrm3 and Chrna3, 4 & 5 are among the top 1.7%, 2.4%, 7.1%, and 8.1% with the strongest positive correlation. **(c)** tPC projection of mouse VISp L1-6, coloured by Chrna4 expression (log CP10K). Upper layer Sst types were selected before PCA to retain the cell type arrangement of Bugeon et al. (Fig. 3f) **(d-g)** Dotplots of cell type-specific cholinergic receptor expression, z-scored across all cells. **(h)** Schematic of RNA count subsampling to control for differences in sequencing depth. Each RNA count from the deeper dataset was sampled with a probability equal to the relative depth of the deep and the shallower dataset (Materials and Methods). **(i-k)** Log2-fold difference in expression with mouse data after subsampling; negative and positive values indicate under- and overexpression, respectively, compared to mouse data. Each violin plot shows the distribution of 100 subsampled datasets. Differences outside of shaded areas are larger than the typical differences between different datasets of the same species (Fig. S12). Without the subsampling procedure, a comparison of raw RNA counts would suggest that every receptor is overexpressed in the mouse by a factor of 8 or more due to the larger sequencing depth of the mouse data. Expression data from refs. [6] (mouse), [9] (VISp L1-3), and [6] (L1-6).

According to our analysis (see Materials and Methods), five cholinergic receptors can predict state modulation of upper-layer subtypes in held-out data in mice (Fig. S11). These receptors also ranked among the top genes in their correlation with state modulation (Fig. 4b). The predictive nicotinic receptors (Chrna3,4,5) showed a rough gradient along tPC1 (see, Fig. 4c). The only predictive inhibitory receptor (Chrm4), on the other hand, was expressed by Pvalb neurons (Figs. 4d), consistent with their negative state modulation (Fig. S1b).

Do the same receptors mediate state modulation in other species? If yes, one would expect differential expression across cell types, with a similar pattern as in mice. However, several receptors that predict state modulation in mice show qualitatively different patterns of expression in the other species (Fig. 4e,f,g). For example, Chrna4 and Chrna5 show much weaker expression in the human data than in mice (Fig. 4e). Chrm4 is overexpressed in the turtle data relative to the other species (Fig. 4f).

The general trend is that the predictive receptors are under-expressed in the other datasets. A possible explanation is a regression to the mean: predictive receptors are, by necessity, expressed in mice. But the relative expression in other datasets could also be due to technical reasons such as a lower sequencing depth (Table 1). Indeed, the typical mouse cell contained several orders of magnitude more RNA counts than the typical human cell (Fig. 4h). We controlled for this confound by downsampling the mouse data to the sequencing depths of the other datasets (Materials and Methods). To measure variability, we also applied this procedure to two datasets from the same species, which revealed typical log2-fold expression differences between -2 and 2 (Fig. S12)—downsampling retained larger differences between species that are qualitatively consistent with the analysis of the full datasets. In the human data, Chrm4, and to a lesser extent Chrna3 & 4, were still underexpressed after downsampling (Fig. 4i). Chrm3 and Chrna5 were underexpressed in the turtle data, whereas Chrm4 was overexpressed (Fig. 4j). In the songbird data, only Chrm4 was overexpressed (Fig. 4k).

Thus, several cholinergic receptors that might mediate state modulation in mice show species-specific expression. This suggests that homologous cell types in different species could show substantial differences in state modulation.

### Robustness of state modulation to cholinergic receptor expression

How do species-specific cholinergic receptor expression patterns influence cortical information flow? Since this depends not just on the cell type-specific gene expression but also on the interplay of different interneurons, we investigated this question using a circuit model (Materials and Methods). We focused on the most salient differences in receptor expression between the three species with a cortex (mice, humans, and turtles).

The model consists of the three most common interneuron types, Pvalb, Sst, and Vip cells, whose connectivity patterns have been mapped [47, 48] and are relatively conserved, at least in mice and humans [49]. Additionally, the computational repertoire of this “canonical circuit” has previously been investigated [50, 51, 52]. The microcircuit connectivity in the turtle (and zebra finch) is, to our knowledge, currently unknown. Our models are therefore a proof-of-principle, to be extended when more data becomes available. As a parsimonious working hypothesis, here we simulated a turtle circuit assuming similar connectivity patterns as in the mammalian cortex and mathematically analysed which changes in connectivity could lead to functional differences (see Network analysis). To explore the effect of cholinergic modulation on excitatory activity, we also included a two-compartmental pyramidal neuron. These two compartments receive different information streams: whereas the soma receives feedforward (sensory) input, the pyramidal dendrites receive top-down input [53, 54, 55]. For visualisation purposes, these input streams were represented by sinusoids of different frequencies (Fig. 5a).

**Figure 5:**
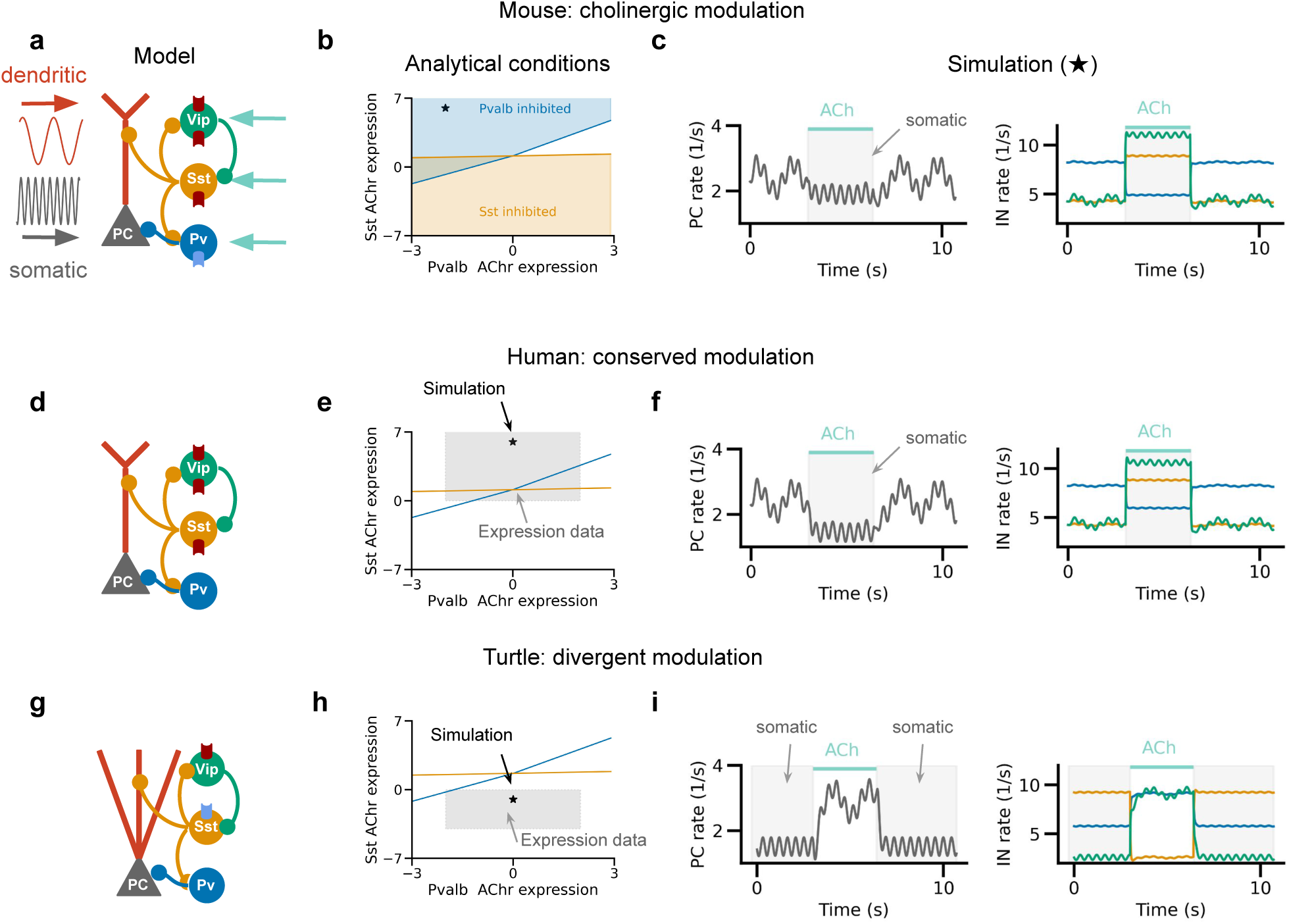
Circuit model predicts functional consequences of cholinergic receptor expression divergence. **(a)** Schematic of mouse cortical circuit model. PC: pyramidal cell. Grey and red sinusoids represent inputs to pyramidal soma and dendrites, respectively; teal arrows represent cholinergic modulation of interneurons. Excitatory and inhibitory ACh receptors are shown in red and blue, respectively. **(b)** Analytical prediction of the Pvalb and Sst ACh receptor expression values for which Pvalb and Sst cells are inhibited by ACh (see Materials and Methods). The star indicates parameter settings used for simulation in **(c)**. At baseline, the PC rate reflects both somatic and dendritic inputs. Cholinergic activation inhibits the dendritic contribution by activating Sst cells. **(d)** Human cortical circuit model, in which Pvalb cells lack inhibitory ACh receptors. **(e)** As (b), but with the shaded area indicating expression values qualitatively consistent with the human data. Star: parameter settings used for simulation in **(f)**. Cholinergic modulation has a qualitatively similar effect due to the indirect cholinergic inhibition of Pvalb neurons via the Sst neurons. **(g)** Turtle circuit model with inhibitory ACh receptor expression in Sst cells. **(h)** As (b), but with the shaded area indicating turtle expression values. Star: parameter settings used for simulation in **(i)**. In contrast to the mammalian circuit models, cholinergic modulation causes a disinhibition of dendritic inputs.

We model cholinergic modulation as an additive input to the interneurons with a strength that is based on cell type-specific receptor expression data (Materials and Methods, Table 4). Cholinergic modulation inhibits the Pvalb population via muscarinic ACh receptors (Chrm4) while activating Vip cells and — to a lesser extent — Sst cells via nicotinic ACh receptors. The activated Sst cells suppress inputs arriving at the dendrites [56, 57, 58], and increase the effective cholinergic inhibition of Pvalb cells. By recruiting dendritic inhibition, ACh therefore limits the influence of top-down inputs on PC rates ([59, 60]; Fig. 5c) while enhancing the influence of top-down inputs [61, 62].

**Table 4:** Network parameters: species and cell type-specific additive cholinergic modulation.

| species \ cell type | Pvalb | Sst | Vip |
| --- | --- | --- | --- |
| Mouse | -2 | 6 | 8 |
| Human | 0 | 6 | 8 |
| Turtle | 0 | -1 | 4 |

How might cholinergic modulation affect interneuron activity and information flow in other species? We modelled a human cortical circuit by deleting the inhibitory Chrm4 receptor from the Pvalb population, mimicking the strongest difference with the mouse VISp data (Fig. 5d). Mathematical analyses indicate that Pvalb activity is still reduced in the presence of ACh due to inhibition from Sst cells (Fig. 5e,f; see Materials and Methods). The difference in direct cholinergic inhibition in Pvalb interneurons between humans and mice might therefore have a relatively weak functional consequence. This is consistent with experimental data [63] and the variable Chrm4 expression across mouse datasets (Fig. S12). By contrast, we found a qualitatively different effect after changing the ACh receptor densities to mimic the turtle data (Fig. 5g): the over-expression of inhibitory Chrm4 receptors by Sst instead of Pvalb cells led to cholinergic disinhibition of dendritic inputs (Fig. 5h,i). This qualitative deviation from the mammalian cortex might affect the state-dependent processing of different input streams in reptiles (see Discussion).

Although the lack of data regarding turtle (or songbird) connectivity precludes us from making testable predictions, these findings highlight that the computation performed by the circuit can be very sensitive to certain patterns of differential expression and robust to others [64, 65].

## Discussion

We have shown that the global gene expression patterns of inhibitory interneurons, as assessed by PCA, show considerable similarity between mice and humans. However, such similarity is not observed between mice and turtles or songbirds. This suggests that the first transcriptomic PC (tPC1) obtained from the upper layers of the mouse cortex [9] may serve as an organizing principle for mammalian interneurons but not for reptilian and avian interneurons. Control analyses indicate that technical factors cannot explain cross-species differences. Instead, our results suggest that the evolutionary divergence of homologous interneurons is mainly explained by changes in gene expression rather than changes in the relative abundance of cell types. Alongside the differences in global expression patterns, we have also observed differences in the expression of cholinergic receptors, suggesting that interneurons undergo species-specific modulation in their functional states.

### Comparison to prior work

The gross transcriptomic differences between species might be surprising given the evolutionary conservation of broad interneuron subclasses in the forebrain [18, 19, 15, 16] and other areas [20, 17]. However, these works also found many genes with species-specific expression, suggesting cell types might be homologous across species but not preserved in their detailed properties. Moreover, fine interneuron subtypes are not necessarily conserved across larger evolutionary distances [33]. Our analyses reveal global differences in gene expression between homologous cell types that rival the differences between glutamatergic cells.

Cholinergic fluctuations with arousal and other cognitive processes have been reported in several species (see, e.g., [66, 67, 68, 69, 70, 71]), and might even be mediated by similar midbrain cell types [72]. However, a causal link between acetylcholine and arousal-like states has only been shown in mice (e.g., [73, 68]), and acetylcholine seems to act via species-specific pathways [74]. For example, most human but not rat PV neurons express the Chrm1 receptor [75]. Even within a single species, cholinergic projections and their effects vary across areas [76, 77, 27] and layers [78, 79]. Although the data therefore suggest a broadly conserved role for acetylcholine in state modulation, this assumption requires testing using functional experiments. In addition, technical differences between datasets naturally pose a more serious limitation for comparing the expression of individual genes than aggregate measures such as PCA. Future work will need to confirm the present results using, e.g., immunohistochemistry.

### Interpretation of transcriptomic PCs

Across all datasets, one feature consistently stands out: the clustering along tPC1&2 by developmental area. A similar pattern has been previously observed based on nonlinear dimensionality reduction and clustering methods (see, e.g., [6, 42]). The structuring of top PCs by developmental origin and cell type is expected since cell types are defined by developmentally-activated transcription factors that coregulate batteries of protein-coding genes [80, 81]. These low-dimensional patterns of gene expression are naturally picked up by a method like PCA.

A clear difference between the mouse datasets is given by two layer-specific subtypes: the deep-layer Meis2 cells [39, 6], and long-range projecting Sst Chodl cells [82, 6]. Their intermediate position along tPC1 (Fig. S9) but distinct connectivity suggests that the correlation of tPC1 with cellular properties [9] might not apply to these deep layer subtypes. This could be further tested using, for example, Patch-seq experiments [83, 84]. A caveat is the relative sensitivity of PCA to cell-type proportions: the intermediate tPC1 scores of Meis2 cells may be caused by their scarcity in the mouse data.

### Evolution of cholinergic modulation

Many genes are at least as predictive of state modulation as cholinergic receptors (Fig. 4b). Some genes might be causally related to state modulation, but others are merely co-regulated with causal genes (Fig. 4a). Thus, strong correlations—the very property that allows the reliable identification of transcriptomic PCs—also preclude the identification of causal genes based on regression analyses. Whether genes predict state modulation also depends on factors only partially under genetic control, such as synaptic connectivity. For example, our network analyses show that synaptic connectivity patterns influence which cholinergic receptor expression differences affect state modulation (Fig. 5): sufficiently strong inhibition from Sst cells causes Pvalb cells to be inhibited by ACh, even if Pvalb cells do not express cholinergic receptors. Finally, species differences in state modulation may not necessarily imply differences in function. An interesting example is turtle Sst cells’ expression of the inhibitory Chrm4 receptor, which might lead to cholinergic disinhibition of pyramidal dendrites. Since sensory inputs to turtle cortex arrive in layer 1 instead of deeper layers [85, 86], we speculate that acetylcholine can thus disinhibit sensory inputs, as it potentially does in mouse cortex. Alternatively, cholinergic modulation could have qualitatively different effects on the processing of somatic and dendritic inputs in turtle compared to mouse cortex. Future experiments could arbitrate between these alternatives.

### Limitations

Our main findings seem robust against potential confounding factors such as sample size and sequencing depth. Yet, computational controls cannot correct all technical differences between the datasets. Some differences may be due to experimental rather than biological factors. Hence, a cross-species comparison using data collected with similar protocols is an important topic for future work. We used theoretical modelling to explore the functional implications of expression differences. At this stage, a lack of experimental constraints prevented us from making specific predictions regarding species-specific effects of internal states. We expect this to change with the availability of more data from non-standard model organisms.

### Conclusion

The wide availability of transcriptomic data in different species offers new opportunities for comparative analyses. Transcriptomic data can not only predict behavioural features such as state modulation but also the electrophysiology and morphology of homologous cell types ([87, 7, 88], but see [24]) which are more accessible. It will be exciting to see whether these predictions generalize across species and if they correlate with high-variance transcriptomic dimensions. More generally, we expect that future cross-species experiments will complement work in genetically accessible mice to reveal general principles of brain function.

## Materials and Methods

Code was written in Python and R and combined into a reproducible workflow using Snakemake [89]. The code will be made available at https://github.com/JoramKeijser/transcriptomic_axes upon publication.

### Datasets

An overview of the analysed datasets is shown in Table 1. The datasets used for each analysis are listed in the figure captions. An overview of the dataset and preprocessing steps is shown in Table2. Table 3 lists the relative frequency of different cell types in each dataset.

### Replication of Bugeon et al

The starting point of our replication was the in vivo calcium imaging data and in situ transcriptomic data previously described by Bugeon et al. [9]. We preprocessed and analysed these data following the original paper unless indicated otherwise. We selected interneurons with a high-confidence assignment to a particular subtype (posterior probability [45] at least 0.5) that belonged to a subtype with at least 3 cells. We used the previous assignment into 35 upper-layer subtypes and grouped Serpinf1 cells into the Vip class. Consistent with the original publication, this resulted in 1065 cells, hierarchically distributed across 5 subclasses (e.g., Pvalb) and 35 subtypes (e.g., Pvalb-Reln-Itm2a). The Npy gene count of 58 cells was missing (NaN); we assumed these values were missing at random and imputed them with the subtype-specific median value. Zero-imputation gave similar results. We computed each cell’s average activity per behavioural state. Whenever a cell was recorded during multiple sessions, we used the session with the longest period of “stationary synchronised” activity since this was the least frequent state. Since 193 cells were not recorded during the stationary synchronised state, state modulation was computed for the remaining 872 cells. The expression matrix contained continuously valued estimates of gene expression instead of integer counts. We normalized these values to 10,000 “counts” per cell for consistency with the other datasets, although this slightly decreased predictive performance. Finally, we log-transformed the normalized values after adding one pseudo-count, log(1 + *x*). The log transformation is a widely used preprocessing step in the analysis of count data [90, 91], although other transformations are also possible (see, e.g., [92, 93]). Linear least squares regression was used to predict state modulation from individual PCs or cholinergic receptors; cross-validated ridge regression was used to predict state modulation from multiple PCs, to mitigate overfitting.

### Other datasets

The transcriptomic datasets each consisted of raw count matrices and metadata that included cell class and subtype/cluster. For the Tasic dataset, we only considered the VISp (not the ALM) cells to allow for a direct comparison with Bugeon et al. In the mouse datasets, we merged the small number of Serpinf1 cells into the Vip cluster for consistency with the analyses from ref. [9]. From the Colquitt and Tosches datasets, we only used the zebra finch and turtle cells, respectively, since the data from other species (Bengalese finch and lizard) contained only a small number of interneurons. For both datasets, we assigned cells to putative mammalian homologues according to the correlation-based matching in the original publications [18, 19]. For the Hodge dataset, we assigned each cell a cell type based on the original publication [14]. Cross-species comparisons were based on one-to-one orthologs, identified using the eggNOG-mapper v2 web interface [94, 95]. We first downloaded the protein fasta files for *Homo sapiens* (GRCh38.p14), *Mus musculus* (GRCm39), *Taeniopygia guttata* (bTaeGut1.4.pri), and *Trachemys scripta elegans* (CAS Tse 1.0). We ran eggNOG-mapper on each file with the Ort. restrictions parameter set to one2one, all other parameters were set to their default values (Min. hit e-value: 0.001; Min. hit bit-score: 60, Min. % of query cov.: 20; Min. % of subject cov.: 20). EggNOG-mapper’s preferred names were used to intersect the results across species, resulting in 12627 one-to-one orthologs, 11002 of which were present in all single cell/nucleus datasets.

### Principal component analysis (PCA)

We scaled gene expression values to 10,000 counts per cell (CP10K) to account for differences in sequencing depth across cells and log-transformed the normalized data. We then identified the top 2000 highly variable genes based on their dispersion across cells (Scanpy’s highly variable genes; using 3000 genes gave similar results). We computed the top 30 PCs based on these highly variable genes. For visual comparison, we made an arbitrary but consistent choice for the signs of tPC1 and tPC2.

We quantified the similarity of PCs from different datasets using principal angles ([96]; Scipy’s subspace angles).

More precisely, let *W_X_* be the gene-by-PC matrix whose columns are the PCs of dataset *A*. The principal angles between the PC subspaces of datasets *A* and *B* are computed from the singular value decomposition (SVD) of the PC-by-PC matrix 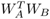, i.e.

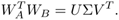

The columns of *U* and *V* contain paired linear combinations of PCs from datasets A and B, respectively, ordered by principal angles. The diagonal matrix Σ contains the singular values *σ_i_*. The *i*th principal angle from the corresponding singular value *σ_i_* is computed as

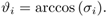

As a complementary measure of PC subspace similarity, we computed the variance explained in one dataset by the top PCs of another dataset. Let **w***_A,i_* be the *i*th PC of dataset *A*, and let *C^B^* be the covariance matrix of dataset *B*. The *i*th PC of dataset *A* explains an amount of variance in dataset *B* equal to

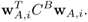

For each pairwise comparison, we computed the covariance and PCs only from genes that were highly variable in both datasets. This was done to avoid the computation of large covariance matrices. For comparison, the variance of each PC was normalized by the variance explained by the first PC of the original dataset:

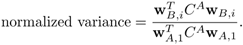

Chance level was estimated by computing the variance explained by a random, normalized vector. To predict state modulation from the tPCs of other species, we first intersected their gene sets with the 72 genes from Bugeon et al. [9] We then separately preprocessed both datasets. Finally, we projected the Bugeon data onto tPC1 from the secondary dataset and used this to predict state modulation. Performance was quantified using leave-one-out *R*^2^ and the Pearson correlation coefficient.

### Subsampling gene counts

The datasets vary in their sequencing depth (the number of RNA counts per cell, see Table 1), presumably due to a combination of technical and biological differences. We aimed to control for these differences by downsampling counts to the depth of the shallower dataset as follows (Figs. 4 S4, S12). Let 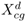 be the number of counts from gene *g* in cell *c* of dataset *d*. We defined the count depth of a dataset as the average counts per cell:

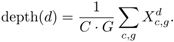

Here *C* is the number of cells, and *G* is the number of genes. If *d*_1_ is the shallowest dataset, and *d*_2_ is a deeper-sequenced dataset, we define their relative sequencing depth as

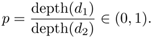

To match the sequencing depth of the shallower dataset, we keep each gene count with a probability *p*:

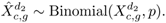

This subsampling procedure equalizes the sequencing depth of the down-sampled dataset to that of the shallower dataset. The Tasic and Hodge data served as the reference datasets for comparison with the mouse and human expression values, respectively, since these were the deepest datasets. The dot plots in Fig. 4 were computed using Scanpy’s dotplot function without the just described downscaling steps. In this case, gene counts were first z-scored to highlight differences between cell types.

### Subsampling cells

To investigate the influence of the number of cells, we downsampled the number of cells in each dataset to that of the smallest dataset (640 cells, Table 1). Sampling cells was done without replacement for 10 different random seeds. After downsampling, we performed preprocessing PCA as described above.

### Dataset integration

We used Seurat’s anchor-based integration [40] to map datasets onto the Tasic data (Fig. S6). To this end, we converted the AnnData objects to Seurat objects. Next, we separately log normalized each dataset as described above (this time using the equivalent Seurat function NormalizeData) and found genes that were highly variable across datasets (FindVariableFeatures, followed by SelectIntegrationFeatures, with 2000 features). Next, we found mutual nearest neighbours across datasets (”anchors”) after projecting each dataset onto the other’s PCA space (reciprocal PCA). A more flexible reduction method (canonical correlation analysis) gave similar results. We then used the anchors to identify and project out dataset-specific differences. After integration, PCA was performed separately on each transformed dataset.

### Correlating gene expression with state modulation

To search across all mouse genes, including cholinergic receptors, we correlated the expression from the Tasic data [6] with the state modulation from the Bugeon data [9]. We first averaged the expression and state modulation within each subtype and then computed the correlation coefficient across subtypes (Fig. 4b). To correlate state modulation from the Bugeon dataset with tPC1 from other datasets (Fig. 2g), we computed each tPC1 based on the 72 genes from the Bugeon et al. in situ panel.

### Network simulations

We simulated a rate-based network of Pvalb, Sst, and Vip interneurons and excitatory pyramidal neurons. A single equation represented each cell type except for the pyramidal neurons, represented by two equations, modelling the somatic and dendritic compartments. The network state was defined by the rate vector **r** = (*r_e_, r_d_, r_p_, r_s_, r_v_*), of somatic, dendritic, Pvalb, Sst, and Vip activity. The rate of cell type/compartment *x* evolved according to

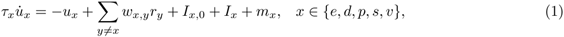

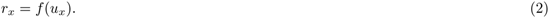

Here *τ_x_* is the membrane time constant (2 ms for excitatory cells, 10 ms for inhibitory cells), *f* (*u*) = max(*u,* 0) is the rectified linear activation function, and the *w_xy_* are recurrent weights. *I_x,_*_0_ is a constant background input that sets the baseline rate, *I_x_* is a time-varying external input, and *m_x_* is an additive cholinergic modulation. We will refer to *m_x_* as a cell’s cholinergic receptor density to distinguish it from the “effective” cholinergic modulation, which also depends on the network dynamics (see Network analysis).

The recurrent connections were chosen based on experimental [47, 49] and theoretical work [52]. The only difference is relatively weak mutual inhibition between Sst and Vip neurons; strong inhibition could prevent the simultaneous activation of these cell types observed in the data [9].

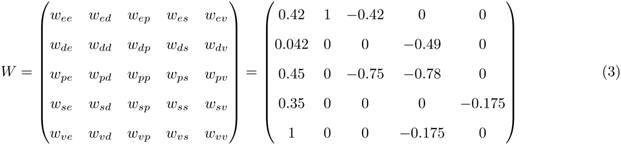

The background inputs *I_x,_*_0_ were set to achieve the following baseline rates:

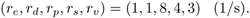

The external inputs to pyramidal soma and dendrites were defined as:

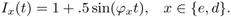

with *φ_e_*= 1*/*300 ms (soma) or *φ_d_*= 1*/*70 ms (dendrite).

The interneurons received cholinergic modulation instead of external inputs, and their amplitudes were varied based on qualitative differences in cholinergic receptor expression (Table 4). These amplitudes were the only differences between species-specific networks. To isolate the effect of differences in ACh receptor densities, the synaptic connectivity patterns were unchanged. In the mouse network, Pvalb neurons were negatively modulated; Vip and — to a lesser extent — Sst neurons were positively modulated. This is consistent with both the activity and expression data from the mouse. For the other species, we only have expression data. In the human and turtle network, Pvalb neurons were not modulated, consistent with their weak or absent expression of, e.g., Chrm4. In the turtle network, Sst neurons were negatively modulated; Vip neurons were positively modulated, but to a smaller extent, given the under-expression of Chrm3 and Chrna5 in the turtle data. For the turtle network, we added a positive external input (amplitude 5) to the Sst equation in the absence of cholinergic modulation. A similar result could be obtained by decreasing the dendritic drive during baseline.

The network dynamics were numerically integrated using a forward Euler scheme with a time step of 0.1 milliseconds. Each simulation consisted of 11000 time steps divided into a baseline period of 3300 steps, a cholinergic modulation time of 3400 steps, and another baseline period of 4300 steps. Not shown in Fig. 5 is an initial settling time of 300 timesteps. These values were chosen to let the figure highlight the effect of turning the modulation on and off.

## Supporting Material

### Replication of Bugeon et al

We validated our analysis pipeline by replicating the relevant results from Bugeon et al. [9] on their data. These data consist of in vivo neural activity and in situ gene expression of neurons from layers 1-3 of mouse primary visual cortex (*Mus musculus* VISp). Expression data was limited to a panel of 72 genes previously selected to identify interneuron subtypes [45]. The data also contain behavioural variables (e.g., running speed) that assign each time point to a “behavioural state”. Bugeon et al. distinguished three possible states: running (distinguished by a positive running speed), stationary desynchronized (zero running speed and little neural oscillations), and stationary synchronised (zero running speed and prominent neural oscillations). A neuron’s state modulation was defined as the normalised difference between its average activity during the most and least active state:

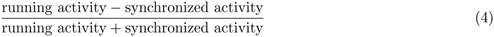

Neurons that are more active during running compared to baseline will therefore have a positive state modulation. In contrast, neurons that are less active during running will have a negative state modulation.

We selected high-quality cells following the criteria from Bugeon et al. ([9], see Materials and Methods), resulting in the same number of 1,065 inhibitory interneurons reported in their work. These interneurons are hierarchically distributed across 5 subclasses and 35 subtypes. State modulation could be computed for 872 neurons recorded during both running and synchronized states. As previously reported, visualising the neural activity during these states suggested differential state modulation between both cell classes and subtypes (Fig. S1a). We then computed each neuron’s state modulation based on its time-averaged activity (Equation (4)). Consistent with ref. [9], state modulation was negative for Pvalb (Pearson correlation -0.13), small and positive for Sst and Lamp5 (both 0.02), and strongly positive for Vip and Sncg interneurons (0.13 and 0.21, respectively) (Fig. S1b).

Next, we aimed to replicate the correlation between the first transcriptomic principal component (tPC1) and the state modulation. To compute tPC1, we first normalised and log-transformed the cell-by-gene expression matrix to correct for differences in sequencing depth and to stabilise the gene-count variances. We then applied principal component analysis to the transformed matrix. State modulation could indeed be predicted from tPC1 projections, both for subtypes (Fig. S1c, leave-one-out *R*^2^ = 0.19) and for individual neurons (*R*^2^ = 0.17). The predictive power of tPC1 is to some extent driven by between-subclass differences in gene expression [37] since it is relatively modest within individual subclasses (Fig. S2). Additional tPCs contained less information regarding state modulation: the second-best tPC (tPC29) achieved an *R*^2^ of 0.10 and explained only 0.9% of the variance, and tPC2 did not predict state modulation at all (*R*^2^ = *−*0.01, Fig. S3). Together, the first 30 tPCs improved upon tPC1 (*R*^2^ = 0.60, 76.2% of total variance).

Finally, we verified the correlation between state modulation and cholinergic receptor expression that might reflect a mechanistic link between state modulation and transcriptome [9]. Since cholinergic receptor expression was not measured for the in vivo recorded neurons (the 72 gene panel did not include these receptors), its relationship with state modulation can only be tested using external expression values. Following ref. [9], we obtained these values from the publicly available data of Tasic et al. [6]. We preprocessed the raw count data like the Bugeon et al. expression matrix and selected the 35 upper-layer subtypes present in the in vivo data. We then computed the average receptor expression of each subtype and compared this with its average state modulation. Linear regression showed that the expression of 5 out of 15 cholinergic receptors (or receptor subunits) could predict state modulation (Figs. S1d, S11). These consist of the 4 receptors shown by Bugeon et al. (Chrm3,4 and Chrna4,5, their Fig. 6b) and an additional nicotinic receptor (Chrna3).

We found one qualitative difference with previous results (Fig. 5c in ref. [9]), namely a clustering of tPC1 scores into two groups corresponding to developmental origin [38]. This was caused by the log transformation used here but not in the original analyses (Fig. S3). The log transformation is a widely used preprocessing step in the analysis of count data [90, 91]. However, other transformations are also possible (see, e.g., [92, 93]). Here, it had only a minor effect on the quantitative relationship between tPC1 and state modulation (Fig. S3).

### Network analysis

The cholinergic receptor densities in our simulations were chosen consistently with the transcriptomic and activity data, but other choices are also possible, of course. We therefore investigated the effect of varying receptor densities using mathematical analyses. In particular, we asked for which receptor densities the cholinergic effect might be different from that in the mouse. For example: does the lack of inhibitory receptors in human Pvalb cells imply that these cells are not inhibited during cholinergic modulation? And does the expression of inhibitory receptors by turtle Sst cells imply that these cells are actually inhibited? In our simulations, all neurons receive net-positive inputs. Under these conditions, the network model contains only one nonlinearity: the rectification of dendritic activity that reaches the soma. The rectification is piecewise linear: if the dendrites are excited, the dendrites influence the soma (*w_ed_* = 1); if the dendrites are inhibited, the dendrite remains inactive and decouple from the soma (*w_ed_* = 0). The network dynamics are, therefore, governed by one of two connectivity matrices that only differ in the entry *w_ed_*. Otherwise, the dynamics are linear:

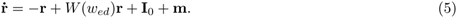

Here, **m** is the vector modelling cholinergic modulation, and **I**_0_ is the external input. For a given somato-dendritic coupling *w_ed_* and cholinergic modulation **m**, the steady state rates are found by solving 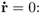:

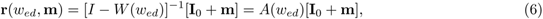

where we defined *A*(*w_ed_*) = [*I − W* (*w_ed_*)]*^−^*^1^ as the matrix that maps inputs to steady-state rates:

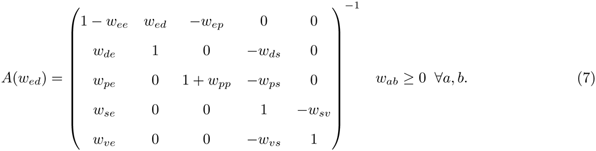

Below, we will compute the relevant entries of *A*(*w_ed_*) up to its determinant, which is positive and therefore does not affect the entries’ signs.

We use equation (6) to compute the modulatory effect on the network activity as the difference between the rates with and without modulation. We will consider the cases in which cholinergic modulation activates the dendrite that was silent without modulation (off*→*on) or inactivates the dendrite that was activated without ACh (on*→*off). The other two cases (on*→*on, off*→*on) can be derived analogously.

First, consider the case that modulation switches the dendrites off, as for the mouse and human circuits.

The resulting change in network activity equals:

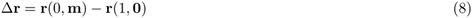

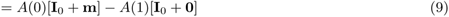

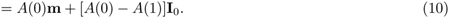

In case that modulation switches the dendrites on, as for the turtle circuit, the resulting change in network activity equals:

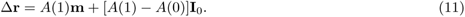

The first term in these equations is a linear combination of the receptor densities **m** = (*m_e_, m_d_, m_p_, m_s_, m_v_*), describing how the cholinergic modulation of individual populations propagates through the network. In our model, pyramidal cells do not express cholinergic receptors (*m_e_* = *m_d_* = 0), such that the cholinergic effect is a linear combination of only the interneuron receptor densities. The second term in Eq. (11) is independent of the precise modulation and describes how the background input **I**_0_ propagates through the network with and without activated dendrites. Since this term is small, we ignore it in the following derivations, but it is shown in Fig. 5.

First, consider the cholinergic effect on Pvalb cells, which equals:

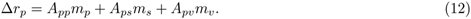

Substituting the entries of *A*(0) gives:

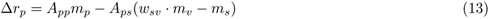

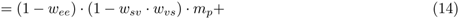

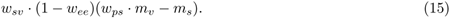

An analogous equation describes Δ*r_s_* (see below). The lines in Figure 5 show Δ*r_p_* = 0 and Δ*r_s_* = 0 as a function of the Pvalb and Sst receptor densities *m_p_* and *m_s_*, for a fixed Vip density *m_v_*. These boundaries delineate domains of positive and negative modulation of Pvalb and Sst interneurons.

So does the absence of inhibitory ACh receptors in human Pvalb cells (Fig. 4; *m_p_* = 0) imply that these cells will not be inhibited? Equation Eq. (13) shows that these cells will still be inhibited indirectly under the condition that:

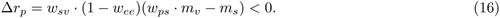

The coefficient *w_sv_ ·* (1 *− w_ee_*) is positive, assuming the recurrent connectivity is not very strong (*w_ee_ ≤* 1). Inhibition of Pvalb cells is then equivalent to

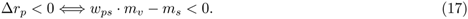

According to the expression data, Sst and Vip cells are positively modulated (*m_s_, m_v_ >* 0). Therefore, Pvalb cells will be inhibited by ACh as long as the inhibition from Ssts is stronger than the disinhibition from Vips.

Under these conditions, the limited Chrm4 expression by human Pvalb cells is compatible with their inhibition.

Let us next consider the differential expression of inhibitory ACh receptors in Sst interneurons in turtles versus mammals. Intuitively, this is expected to cause a cholinergic suppression of Sst cells in the turtle, in contrast to the mouse. In the model, the cholinergic effect on Sst cells equals:

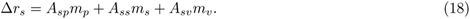

Subsituting the entries from *A*(1) gives:

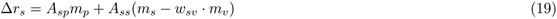

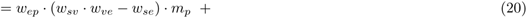

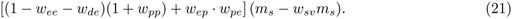

For the turtle circuit, *m_p_ ≤* 0, since Pvalb cells (weakly) express inhibitory ACh receptors. The first term will therefore be negative if *w_sv_ · w_ve_ − w_se_ >* 0. Further, *m_s_ <* 0 and *m_v_ >* 0, such that *m_s_ − w_sv_m_v_ <* 0. The contribution of the second term will therefore be negative if:

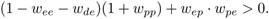

This will be the case unless recurrent excitation is very strong or the feedback loop between PCs and Pvalb cells is very weak. In summary, the expression of inhibitory ACh receptors by turtle Sst cells will indeed lead to their cholinergic inhibition, provided that the excitation onto Sst cells and the recurrent excitation are not too strong.

### Software

An Anaconda [97] environment with the appropriate software will be provided along with the code (Table 5).

**Table 5:** Software versions.

| Software | version |
| --- | --- |
| AnnData [98] | 0.8.0 |
| Matplotlib [99] | 3.6.2 |
| Numpy [100] | 1.23.5 |
| Pandas [101] | 1.5.2 |
| Python [102] | 3.10.10 |
| R [103] | 4.3.0 |
| Scanpy [104] | 1.9.1 |
| Scikit-learn [105] | 1.2.1 |
| Scipy [106] | 1.9.3 |
| Seaborn [107] | 0.12.2 |
| Seurat [108] | 4.0 |
| Snakemake [89] | 7.8.2 |
| Statsmodels [109] | 0.13.5 |

## Supplementary figures

**Fig. S1:**
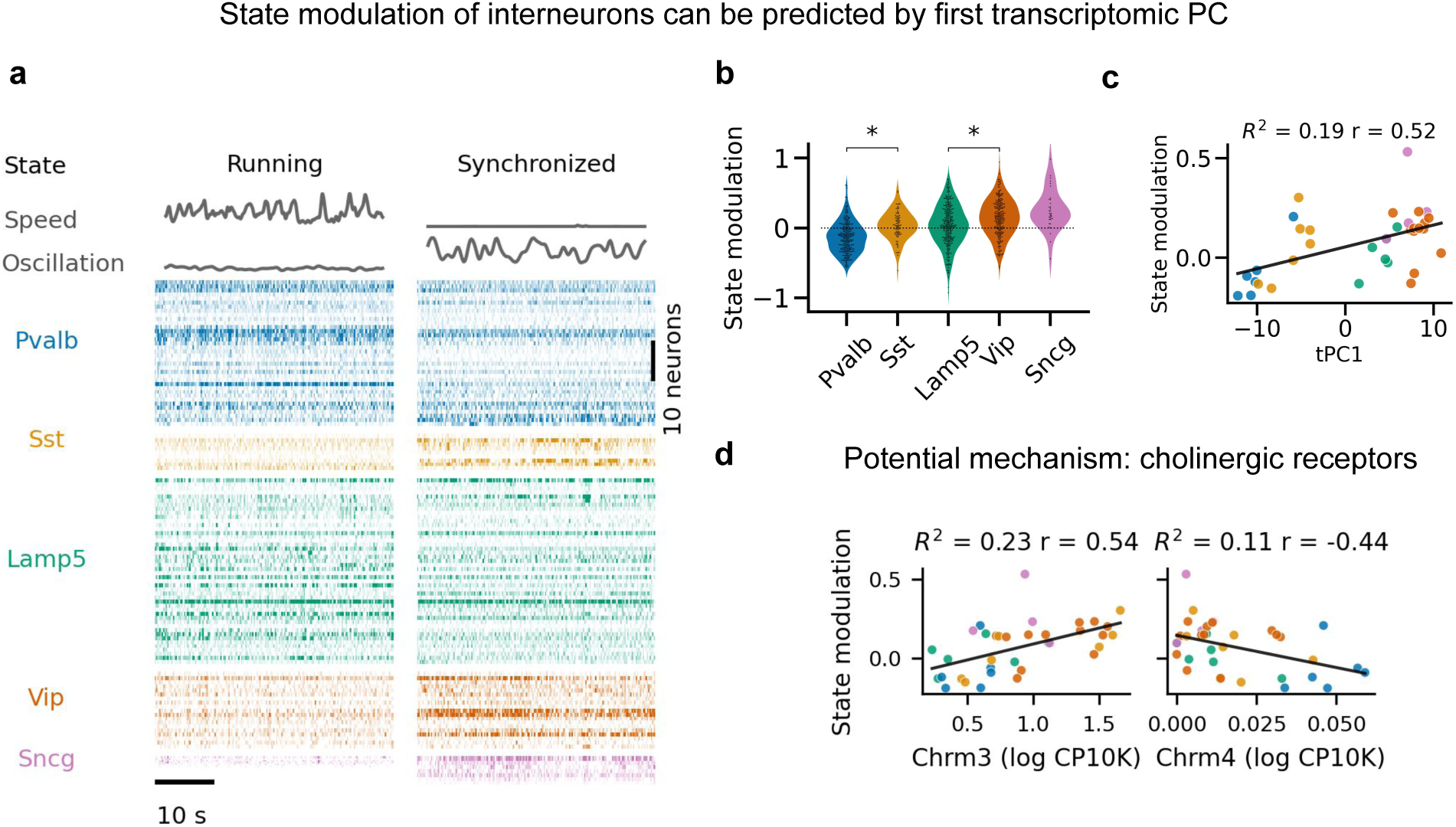
Replication of previous findings from Bugeon et al. [9] **(a)** Neural activity systematically varies with behavioural state (measured by running speed and neural oscillations, see Materials and Methods) both between and within interneuron classes of mouse primary visual cortex (VISp) L1-3. **(b)** State modulation across all sessions for *n* = 872 interneurons. Stars indicate statistically significant differences between subclasses (*p <* 0.05, Mann-Whitney U test). **(c)** The first transcriptomic principal component (tPC1) of the cell-by-gene matrix predicts state modulation of subtypes (*n* = 35); Fig. S2a shows the relationship for individual cells. *R*^2^, leave-one-out fraction of variance explained; *r*, Pearson correlation. Note the two clusters along tPC1, consisting of MGE-derived (Pvalb & Sst) and CGE-derived (Lamp5, Vip, Sncg) interneurons. **(d)** Cholinergic receptors potentially link a neuron’s transcriptome and state modulation. For example, interneurons that overexpress the excitatory receptor Chrm3 are positively state-modulated (*r* = 0.54; *p* = 0.0008), those that overexpress the inhibitory cholinergic receptor Chrm4 are negatively state-modulated (*r* = *−*0.44, *p* = 0.0075). CP10K, counts per 10 thousand. Data and findings from Bugeon et al. [9]. Cholinergic receptor expression in (d) from Tasic et al. [6].

**Fig. S2:**
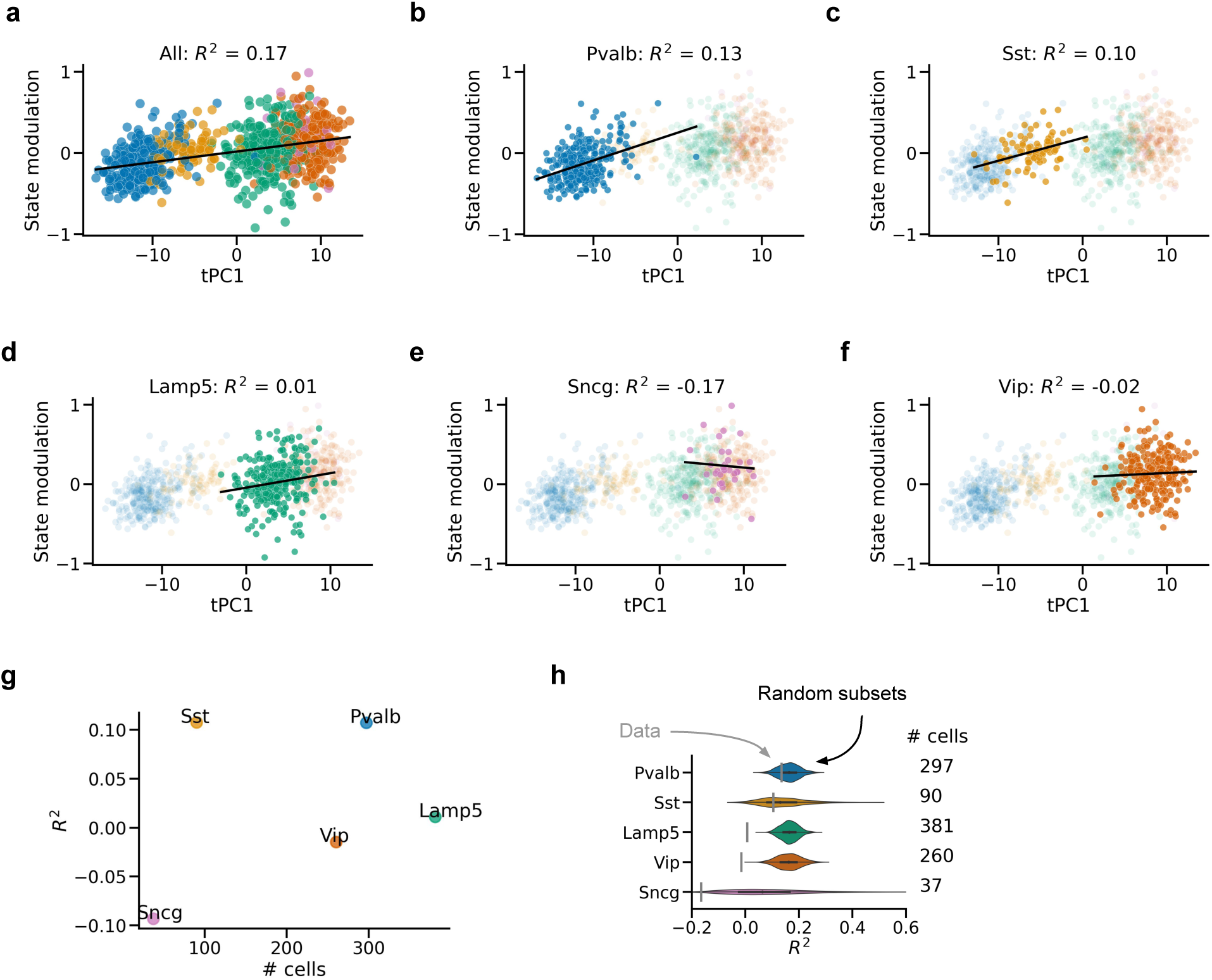
Predicting state modulation from tPC1. Regression based on all cells (**a**) or cells from a single subclass (**b-f**). Predictive performance is worse for individual classes and only better than chance for Pvalb and Sst cells. The correlation between tPC1 and state modulation is therefore partially driven by between-subclass differences. However, tPC1 is still predictive of state modulation across all cells while controlling for subclass (*p* = 0.003, linear mixed model with subclass as random effect). *R*^2^: leave-one-cell-out fraction of variance explained; *R*^2^ *<* 0 indicates a worse fit compared to predicting the same state modulation for each cell independent of tPC1 score. **(g,h)** Poor performance for certain subclasses is not due to a smaller sample size. **(g)** Sample size is not correlated with worse performance. **(h)** Size-matched subsets of all cells outperform below-chance subclasses, except for Sst cells. Grey bars: *R*^2^ values for each subclass. Violin plots: distribution of *R*^2^ values for 1000 random subsets of all cells with sample size matched to the subclass. Data from Bugeon et al. [9].

**Fig. S3:**
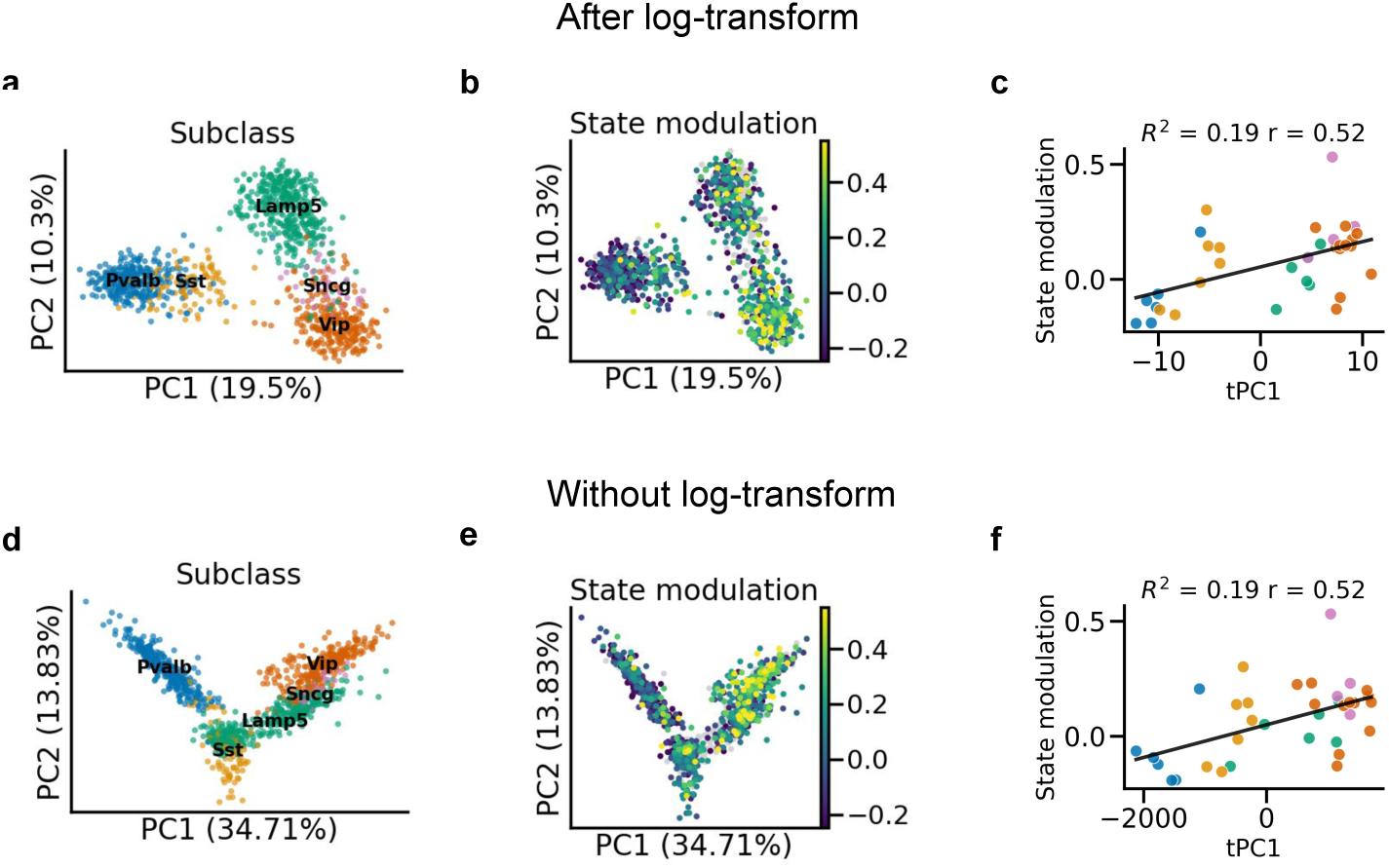
Log-transformation leads to clustering by developmental origin. **(a)** First 2 transcriptomic principal components (tPCs) of the log-transformed count RNA data. **(b)** As (a), with colour indicating state modulation. **(c)** The first transcriptomic PC (tPC1) of log-transformed data predicts state modulation, replicated from Fig. S1c for comparison. *R*^2^: leave-one-out fraction of variance explained, *r*: Pearson correlation. **(d-f)** As (a-c), but without log-transformation. Interneurons now form a continuum along tPC1, but the quantitative relationship between tPC1 and state modulation is preserved (up to 2 digits). Data from Bugeon et al. [9].

**Fig. S4:**
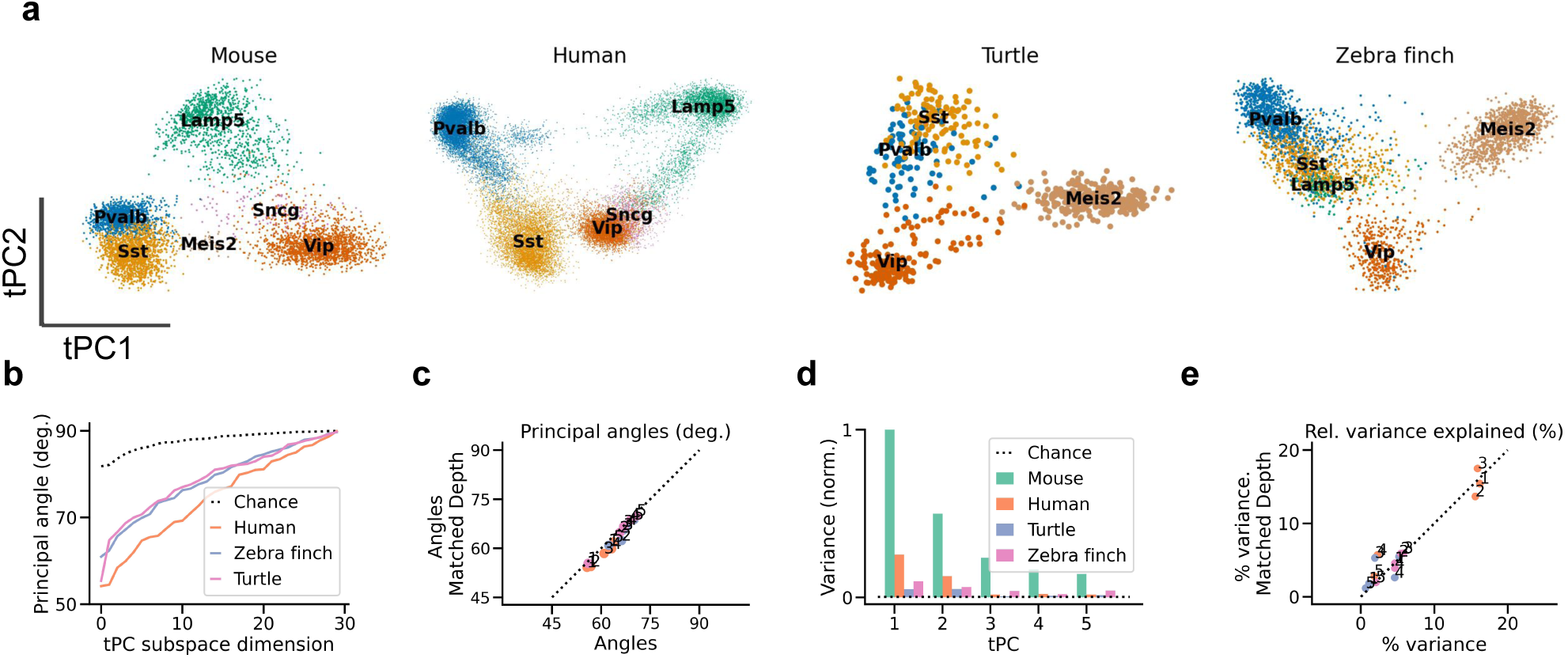
Transcriptomic PCs robust to sequencing depth. **(a)** Projection of each dataset onto its first 2 transcriptomic PCs, after subsampling gene counts to the depth of the shallowest dataset (zebra finch, see Table 1). **(b)** Principal angles between tPC subspaces of subsampled data. **(c)** Comparison between angles of full-depth data and subsampled data. **(d,e)** As (b,c) but for variance explained. The human, turtle, and zebra finch tPC1 explain 15.4%, 1.8%, and 4.6% of the variance explained by mouse tPC1, respectively. Data from refs. [6] (mouse), [15] (human), [18] (turtle), [19] (zebra finch).

**Fig. S5:**
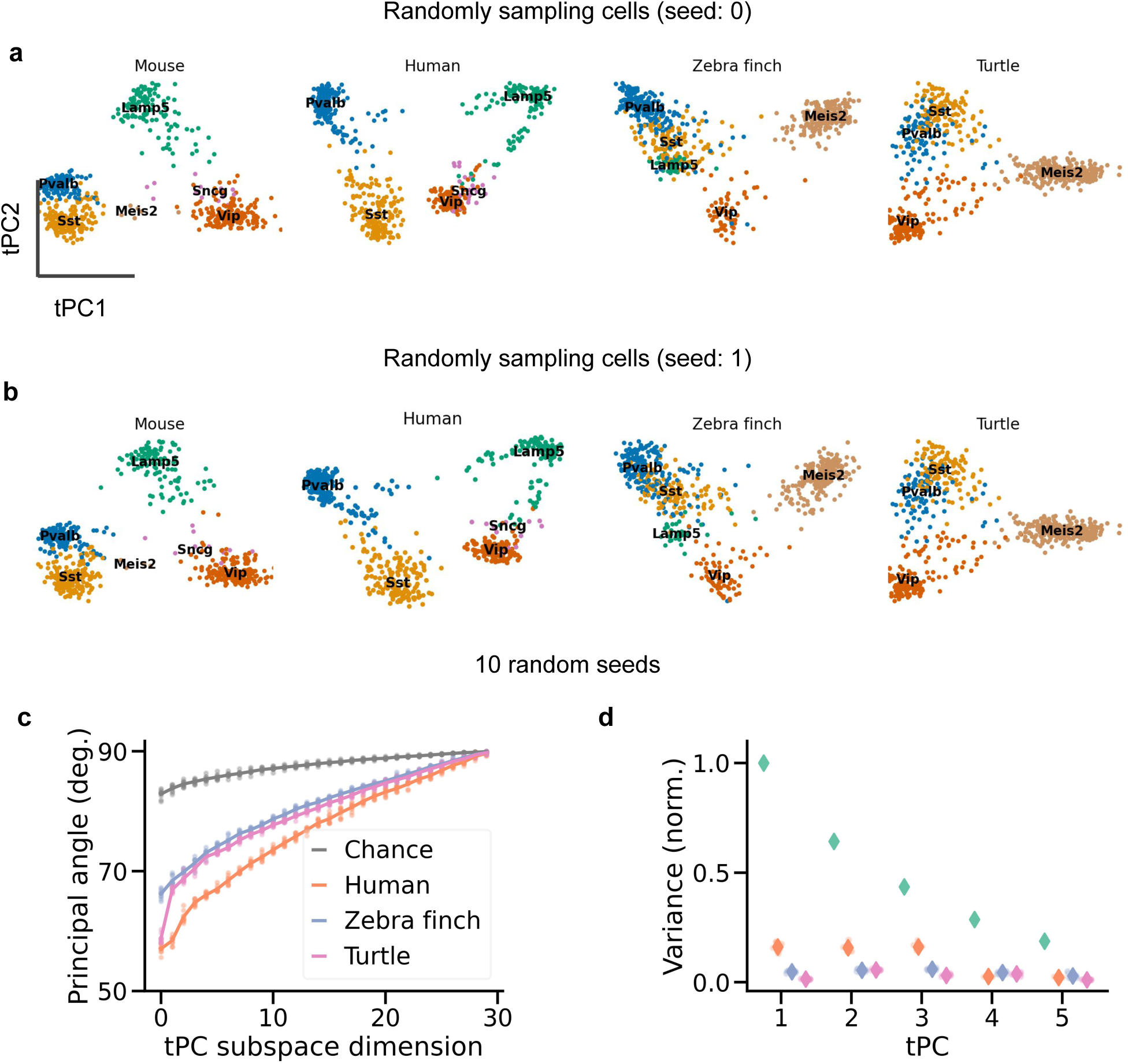
Differences in tPCs not due to cell count. **(a)** Projection of each dataset onto its first 2 transcriptomic PCs, after random sampling of 640 cells (the number of cells in the smallest dataset, with Turtle cells) without replacement. **(b)** Like (a), but for a different random seed. Note that the Turtle plots in (a) and (b) are the same because there is only one way to sample exhaustively without replacement. **(c)** Principal angles between tPC subspaces. Lines: average across 10 random seeds; dots: individual seeds. **(d)** As (c) but for variance explained, normalized by the variance explained by tPC1 from the mouse data. Diamonds: average across 10 random seeds; dots: individual seeds (largely invisible). Colors as in (c); green dots correspond to the mouse data. Data from refs. [6] (mouse), [15] (human), [18] (turtle), [19] (zebra finch).

**Fig. S6:**
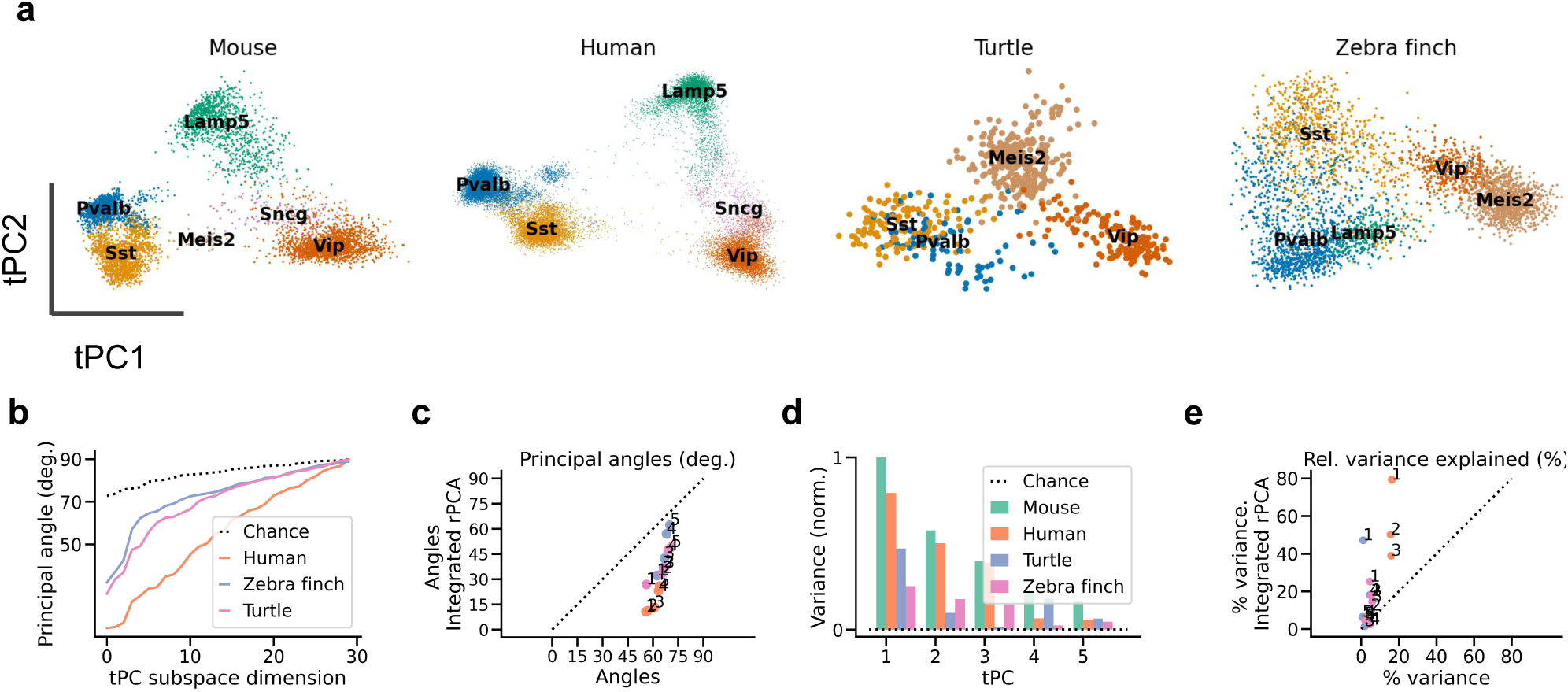
Computational integration increases similarity of mouse and human data. **(a)** Projection of each dataset onto its first 2 transcriptomic PCs, after computational integration. Mouse and human datasets show increased similarity, but turtle cells no longer cluster by cell type (colour). **(b)** Principal angles between tPC subspaces computed after integration. **(c)** Comparison between angles computed without integration. Integration increased the similarity of all datasets, especially of the human data. **(d,e)** As (b,c) but for variance explained. The human, turtle, and zebra finch tPC1 explain 79.4%, 47.2%, and 25.2% of the variance explained by mouse tPC1, respectively. Data from refs. [6] (mouse), [15] (human), [18] (turtle), [19] (zebra finch).

**Fig. S7:**
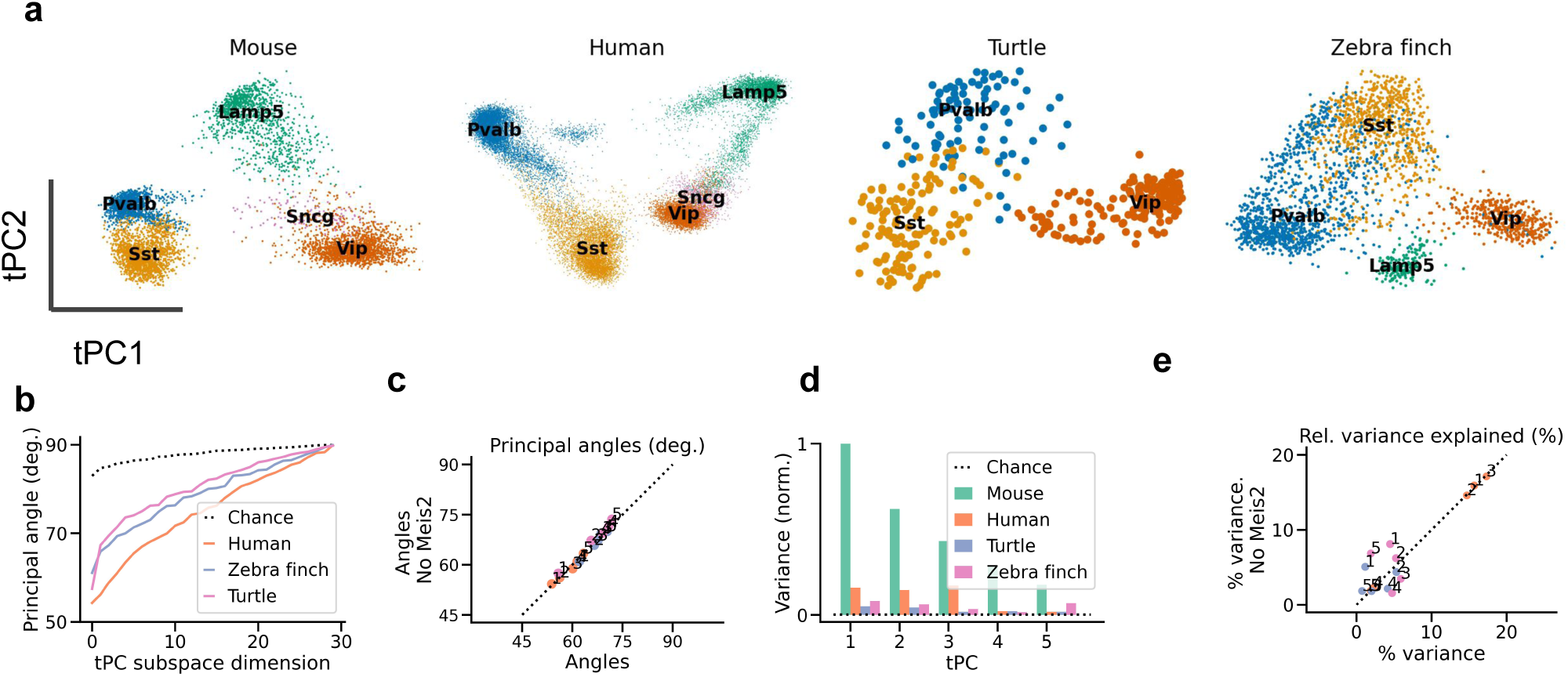
Differences in tPCs not due to Meis2 cells. **(a)** Projection of each dataset onto its first 2 transcriptomic PCs, after removing Meis2 cells. **(b)** Principal angles between tPC subspaces computed without Meis2 cells. **(c)** Comparison between angles computed on all cells vs. cells without Meis2 population. **(d,e)** As (b,c) but for variance explained. The human, turtle, and zebra finch tPC1 explain 16.4%, 5.0% and 8.6% of the variance explained by mouse tPC1, respectively. Data from refs. [6] (mouse), [15] (human), [18] (turtle), [19] (zebra finch).

**Fig. S8:**
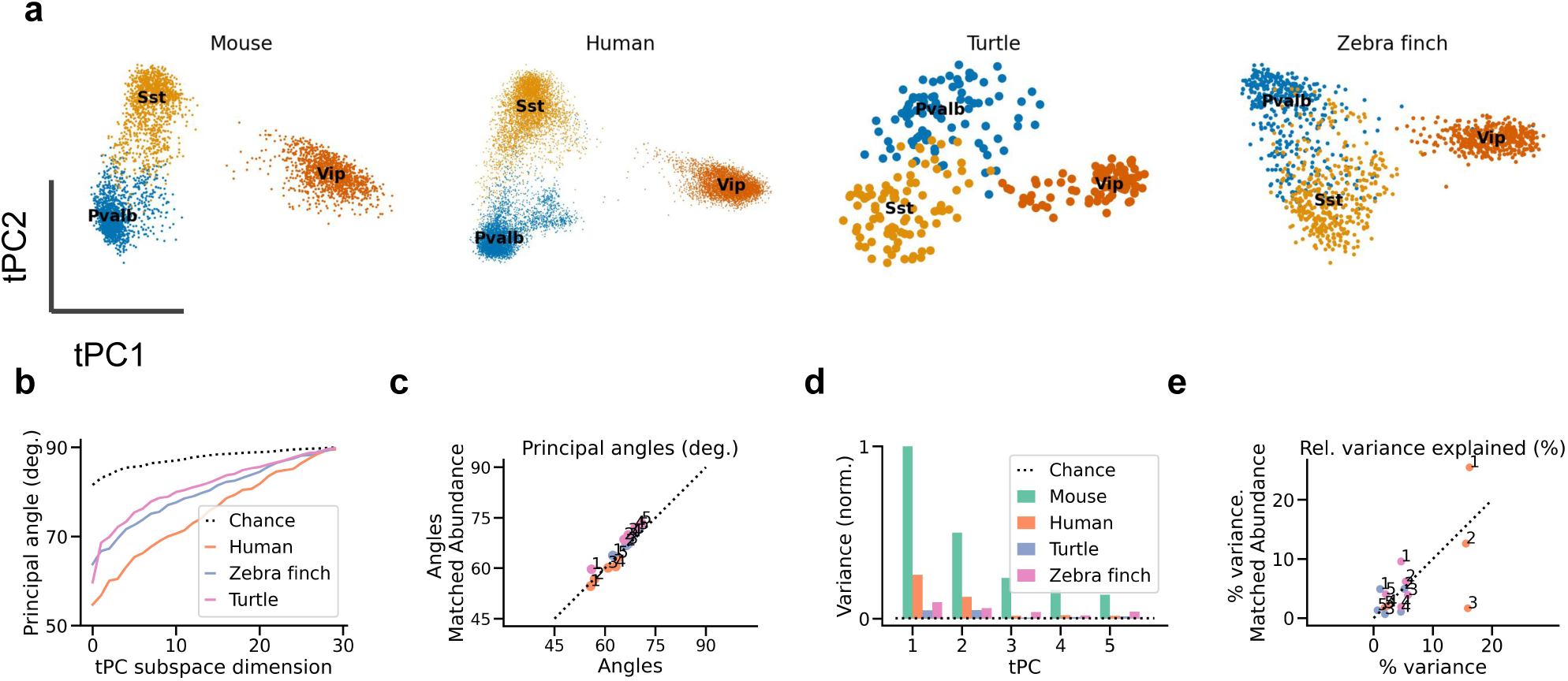
Differences in tPCs not only due to cell type abundance. **(a)** Projection of each dataset onto its first 2 transcriptomic PCs, after matching cell type abundances (Fig. 2h). **(b)** Principal angles between tPC subspaces. **(c)** Comparison between angles computed on all cells vs. cells after matching frequencies. **(d,e)** As (b,c) but for variance explained. The human, turtle, and zebra finch tPC1 explain 25.5%, 4.9%, and 9.5% of the variance explained by mouse tPC1, respectively. Data from refs. [6] (mouse), [15] (human), [18] (turtle), [19] (zebra finch).

**Fig. S9:**
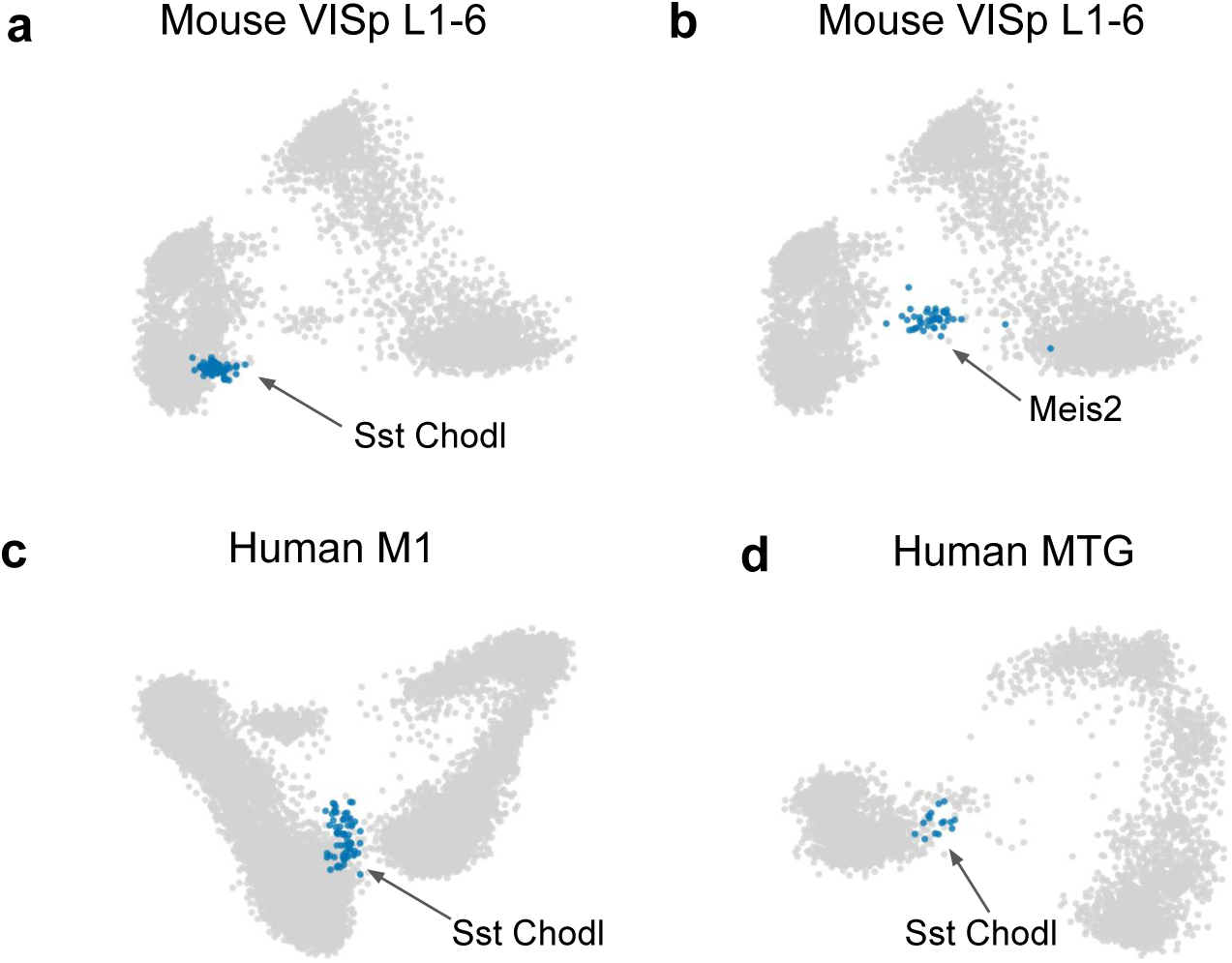
Intermediate tPC1 position of Chodl and Meis2 neurons. Long range projecting Sst-Chodl **(a)** and white matter Meis2-Adamts19 cells **(b)** occupy an intermedate position along tPC1. **(c,d)** Sst Chodl neurons also have intermediate tPC1 scores in the human data. The human datasets do not contain Meis2 cells. Data from refs. [6] (a,b), [15] (c), and [14] (d).

**Fig. S10:**
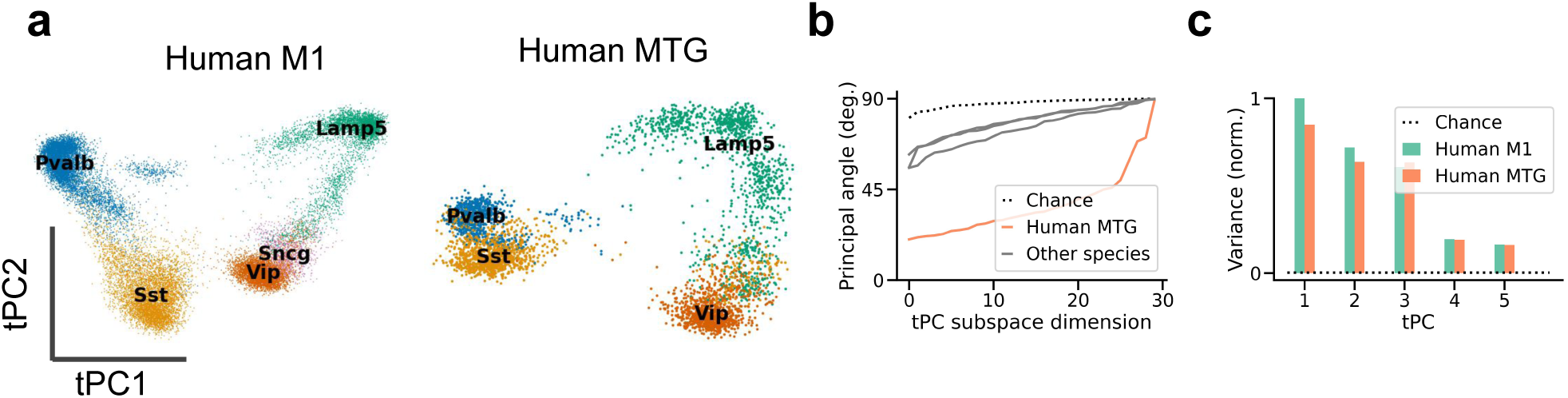
Small differences in PCs of human datasets. **(a)** Projection of human datasets onto their first 2 tPCs. **(b)** Quantification of tPC similarity using principal angles between tPC subspaces of M1 and MTG data. **(c)** Quantification by variance explained in MTG data. The M1 tPC1 explains 85% of the MTG variance explained by the MTG tPC1. Data from refs. [15] and [14].

**Fig. S11:**
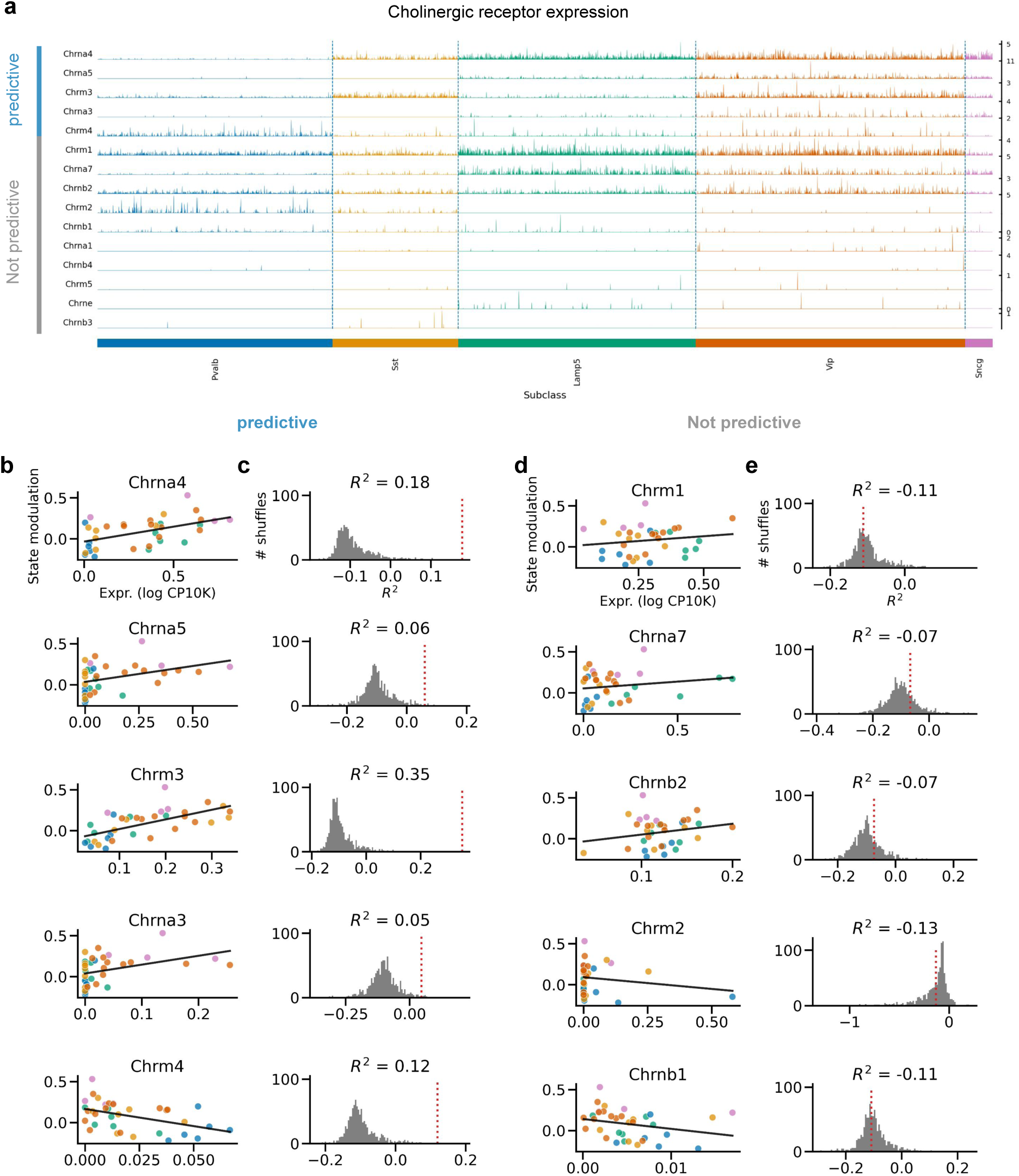
Predicting state modulation from cholinergic receptor expression. **(a)** Tracks plot of cholinergic receptor (subunit) expression. The first 5 receptors predict state modulation; the remaining 10 do not (see b-e). Predictive and unpredictable receptors are independently sorted by expression based on expression levels. Shown are all receptors with an expression of at least 1 count per 10K. **(b)** Relationship between state modulation and log expression of receptors that are predictive of state modulation (1000 permutations, *p <* 0.05). **(c)** Grey: Null distribution of leave-one-out *R*^2^ estimated by linear regression after permuting expression levels. Red: *R*^2^ without permutation. **(d,e)** As (b,c) but for the 5 unpredictive receptors with the highest expression. Receptor expression from Tasic et al. [6]; state modulation from Bugeon et al. [9].

**Fig. S12:**
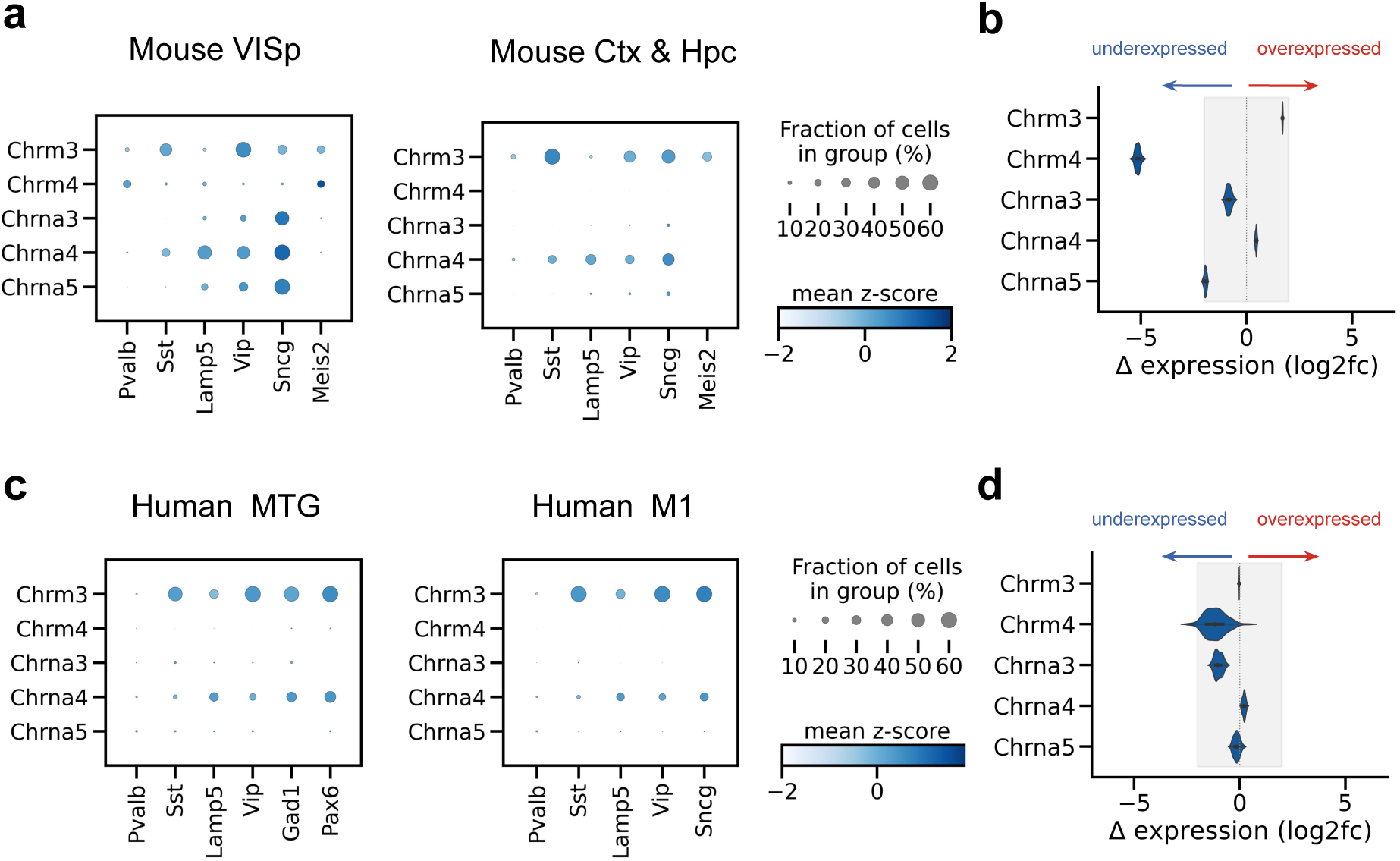
Mostly small within-species differences in ACh receptor expression. **(a)** Dot plots showing the expression of the cholinergic receptors that predict state modulation in mouse VISp L1-3 **(b)** Log2-fold differences in expression after downsampling the VISp dataset to equal sequencing depth as the Ctx & Hpc data. Shaded area: log-fold difference of *±*2, the range of most within-species differences. The exception is Chrm4, which is underexpressed in the Ctx & Hpc data compared to the VISp data. **(c,d)** As (a,b), but for human datasets. The MTG dataset was downsampled to match the M1 data. Data from refs. [6] (mouse VISp), [29] (mouse Ctx & Hc), [14] (human MTG), and [15] (human M1).

